# 15-Lipoxygenase-Mediated Lipid Peroxidation Regulates LRRK2 Kinase Activity

**DOI:** 10.1101/2024.06.12.598654

**Authors:** Matthew T. Keeney, Eric K. Hoffman, Julie Weir, Weston G. Wagner, Emily M. Rocha, Sandra Castro, Kyle Farmer, Marco Fazzari, Roberto Di Maio, Andrei Konradi, Teresa G. Hastings, Sean A. Pintchovski, William D. Shrader, J. Timothy Greenamyre

**Author notes:** **Corresponding author:** J. Timothy Greenamyre University of Pittsburgh 3501 Fifth Avenue, Suite 7039 Pittsburgh, PA 15260. **One Sentence Summary**: 15-Lipoxygenase-derived 4-HNE stimulates LRRK2 kinase activity.

## Abstract

Mutations in *leucine-rich repeat kinase 2 (LRRK2)* that increase its kinase activity are strongly linked to genetic forms of Parkinson’s disease (PD). However, the regulation of endogenous wild-type (WT) LRRK2 kinase activity remains poorly understood, despite its frequent elevation in idiopathic PD (iPD) patients. Various stressors such as mitochondrial dysfunction, lysosomal dyshomeostasis, or vesicle trafficking deficits can activate WT LRRK2 kinase, but the specific molecular mechanisms are not fully understood. We found that the production of 4-hydroxynonenal (4-HNE), a lipid hydroperoxidation end-product, is a common biochemical response to these diverse stimuli. 4-HNE forms post-translational adducts with Cys2024 and Cys2025 in the kinase activation loop of WT LRRK2, significantly increasing its kinase activity. Additionally, we discovered that the 4-HNE responsible for regulating LRRK2 is generated by the action of 15-lipoxygenase (15-LO), making 15-LO an upstream regulator of the pathogenic hyperactivation of LRRK2 kinase activity. Pharmacological inhibition or genetic ablation of 15-LO prevents 4-HNE post-translational modification of LRRK2 kinase and its subsequent pathogenic hyperactivation. Therefore, 15-LO inhibitors, or methods to lower 4-HNE levels, or the targeting of Cys2024/2025 could provide new therapeutic strategies to modulate LRRK2 kinase activity and treat PD.

## INTRODUCTION

Missense mutations in *leucine-rich repeat kinase 2* (LRRK2) are a common cause of familial Parkinson’s disease (PD) (*1, 2*). Both coding and non-coding variants in *LRRK2* have been shown to increase the risk of developing PD (*3, 4*). Most of these mutations are located in the Roc-Cor GTPase or kinase domain of LRRK2 and often result in elevated kinase activity, leading to a toxic gain-of-function (*1, 2, 5, 6*). Evidence also suggests wild-type (WT) LRRK2 kinase activity is enhanced in idiopathic PD (iPD) (*7–11*). Thus, LRRK2 might be a link between genetic and idiopathic forms of PD. However, the upstream signaling mechanisms that promote WT LRRK2 kinase activation are not well understood.

LRRK2 uses a classical kinase switch activation mechanism (*12*), where the dynamics of the kinase domain drive its activation (*13*). Upon activation, LRRK2 phosphorylates its substrates, including a subset of Rab GTPases (e.g., Rab10 and Rab12) (*14, 15*), which regulate endolysosomal function (*8*). Excessive LRRK2 kinase activity can also cause endolysosomal dysfunction, creating a complex feedback loop.

The kinase domain of LRRK2 is similar to the redox-sensitive MAP kinase, ASK1 (*16*), which undergoes redox-induced functional changes. Redox stress, long implicated in PD pathogenesis, can induce WT LRRK2 kinase activity (*5, 9, 17, 18*), potentially to the same extent as the kinase activating G2019S mutation (*9, 18*). Interestingly, LRRK2 can be activated by various stimuli that induce oxidative damage, lysosomal damage, membrane rupture or remodeling, vesicular trafficking deficits, and signaling through Stimulator of Interferon Genes (STING) (*19–24*).

A potential common factor among these stimuli is their association with membrane damage and lipid peroxidation, leading to the accumulation of 4-hydroxynonenal (4-HNE), a reactive aldehyde that plays a role in activating other kinases (*25–28*). The formation of 4-HNE occurs either nonenzymatically when lipids react with reactive oxygen species (ROS), or enzymatically through the hydroperoxidation of membrane phospholipids by lipoxygenase enzymes, primarily 15-lipoxygenase (15-LO) (*29, 30*). 4-HNE can form irreversible Michael-adducts with cysteine residues, and the kinase activation loop of LRRK2 contains highly conserved, solvent-exposed, vicinal cysteine residues, Cys2024 and Cys2025, which may be important for redox sensing (*12*).

Here, we provide evidence that oxidative stimulation of endogenous LRRK2 kinase activity by H_2_O_2_, rotenone, chloroquine, or monensin depends on cysteine residues, Cys2024 and Cys2025, and involves the formation of adducts between 4-HNE and LRRK2. Importantly, blocking 4-HNE production with a 15-LO inhibitor, or through genetic ablation of 15-LO, prevents pathologically elevated LRRK2 kinase activation without affecting basal LRRK2 kinase activity.

## RESULTS

### Stimulated endogenous LRRK2 activity is redox dependent

The regulation of WT LRRK2 kinase activity remains poorly understood, but evidence suggests it is sensitive to redox stress (*8*). Several reports indicate that LRRK2 kinase activity increases with H_2_O_2_ and decreases with the antioxidant curcumin (*9, 18, 31, 32*). WT endogenous LRRK2 kinase activity can be elevated dose-dependently by physiological concentrations of H_2_O_2_, and this can be blocked by α-tocopherol (*9*), which is both an antioxidant and a 15-LO inhibitor (*33*). To further investigate this regulation, we examined the *in situ* kinetics of H_2_O_2_-induced endogenous WT LRRK2 activation in human embryonic kidney 293 cells (HEK293). Cells treated with 5μM H_2_O_2_ for 5, 10, and 15 minutes showed a robust increase in LRRK2 kinase activity, as detected by proximity ligation (PL) assay for the phosphorylation status of the autophosphorylation site Ser1292 (PL pS1292–LRRK2) (*9, 34*). This signal was maximal at 10 minutes and sustained for at least 30 minutes (***Fig. 1A, B***). Pretreatment of cells with a thiol antioxidant, N-acetylcysteine (NAC), prevented the H_2_O_2-_induced increase in PL pS1292-LRRK2 signal (p<0.0001; two-way analysis of variance (ANOVA) with Tukey correction) (***Fig, 1C, D***), confirming the redox sensitivity of LRRK2 kinase activation.

**Figure 1.**
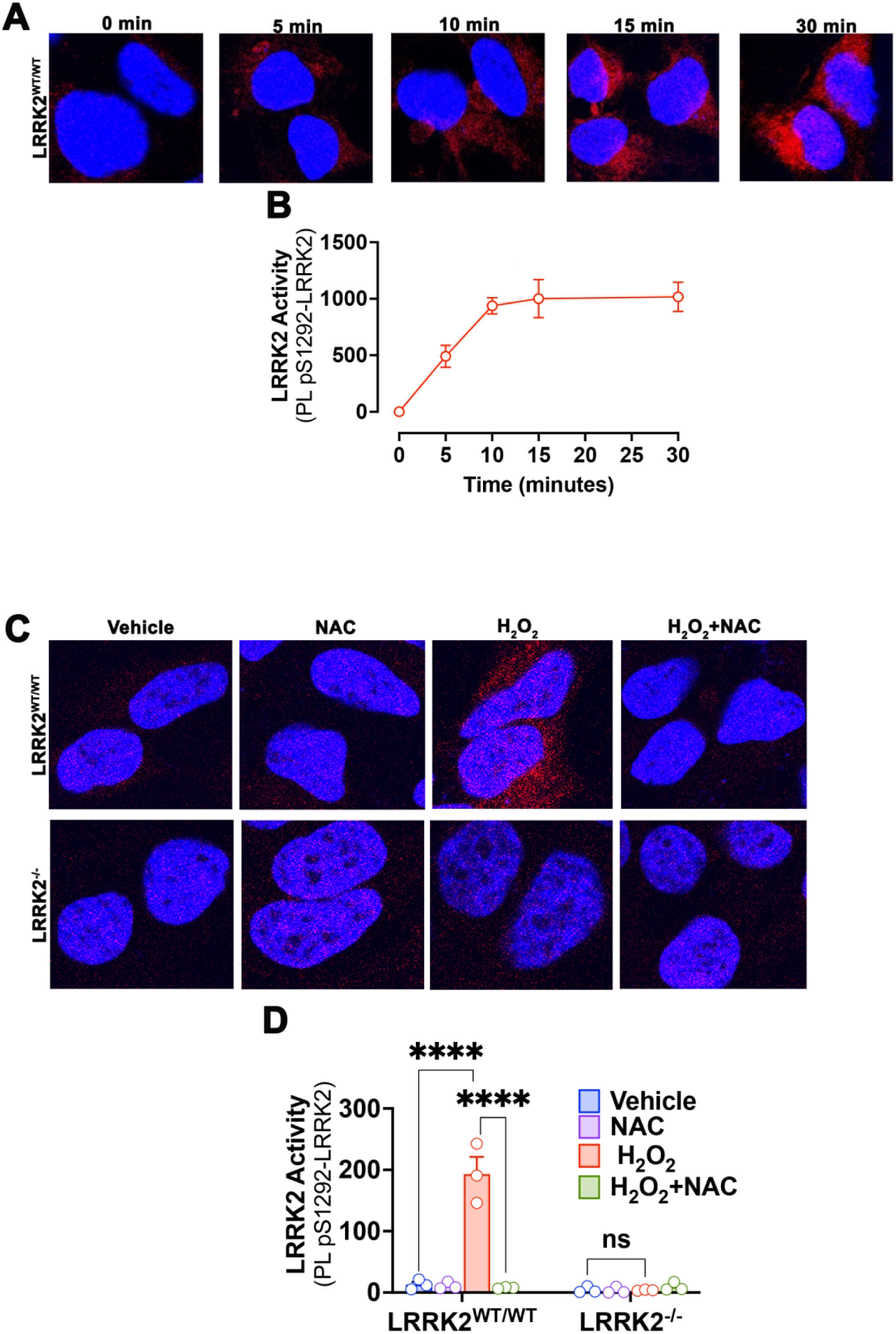
H_2_O_2_ elicits rapid LRRK2 autophosphorylation. (**A**) Endogenous LRRK2 kinase activity was assessed by proximity ligation (PL) assay between pS1292 and total LRRK2 (PL pS1292–LRRK2) in LRRK2^WT/WT^ HEK293 cells treated with 5μM H_2_O_2_ for 5, 10, 15, or 30 minutes. A robust increase in PL pS1292-LRRK2 signal (red) was observed within 10 minutes of treatment and remained sustained. (**B**) Quantification of the time course of LRRK2 activity in response to H_2_O_2_. Each symbol represents the average PL pS1292-LRRK2 fluorescence intensity from 4 independent experiments obtained from 100-150 cells per treatment group for each independent experiment. (**C**) H_2_O_2_ (5μM) induced LRRK2 kinase activity was assessed by PL pS1292-LRRK2 in LRRK2^WT/WT^ and LRRK2^-/-^ HEK293 cells. H_2_O_2_ caused a significant increase in PL pS1292-LRRK2 signal in LRRK2^WT/WT^ compared to vehicle. Pretreatment with NAC prevented the H_2_O_2-_induced increase in PL pS1292-LRRK2 signal, and no signal was detected in LRRK2^-/-^ cells. (**D**) Quantification of LRRK2 activity by PL fluorescence signal. Each symbol represents the average PL pS1292–LRRK2 fluorescence intensity obtained from 100-150 cells per treatment group in three independent experiments. Statistical testing by two-way ANOVA with Tukey correction; ****denotes p<0.0001; ns denotes not significant.

Activation of WT LRRK2 occurs in response to various cellular stressors, including mitochondrial dysfunction, lysosomal dyshomeostasis, and vesicular trafficking deficits (*9, 20–22, 35*). We explored whether known LRRK2 activators such as rotenone, chloroquine, and monensin activate LRRK2 via a redox mechanism. Rotenone, a mitochondrial complex I inhibitor, increases ROS production, redox stress, and mitochondrial dysfunction in cells (*36*). In WT HEK293 cells, sublethal rotenone treatment elicited an increase in PL pS1292-LRRK2 signal compared to vehicle (p<0.0001; two-way ANOVA with Tukey correction) (***Fig. 2A, B***), and this increase was prevented by co-treatment with NAC (p<0.0001; two-way ANOVA with Tukey correction) (***Fig. 2A, B***). Congruent results were obtained by western blotting for the phosphorylation state of the LRRK2 substrate, Rab10 (*14*) (***fig. S1A, B)***. Rotenone treatment increased pT73-Rab10 levels compared to vehicle (p<0.05; two-way ANOVA with Tukey correction), and this increase was prevented by NAC co-treatment (p<0.05; one-way ANOVA with Tukey correction) (***fig. S1A, B***). We confirmed these findings using a third method, a PL assay to amplify the specific signal of pT73-Rab10: PL pT73(Rab10)-Rab10 ***(fig. S2)***. Using this assay, there was a baseline pT73(Rab10)-Rab10 signal in WT HEK293 cells that was reduced in the presence of the LRRK2 kinase inhibitor, PF360 (p<0.0001; two-way ANOVA with Tukey correction) (***fig S2A, B***). Importantly, the baseline pT73(Rab10)-Rab10 signal in WT HEK293 cells was higher than that in LRRK2^-/-^ HEK293 cells (p<0.0001; two-way ANOVA with Tukey correction) (***fig S2A, B).*** Similar to the results obtained with PL pS1292-LRRK2 and pT73-Rab10 western blotting, rotenone elicited an increase in PL pT73(Rab10)-Rab10 signal (p<0.0005; two-way ANOVA with Tukey correction), that was prevented by co-treatment with NAC (p<0.0001; two-way ANOVA with Tukey correction) (***fig. S1D, E***).

**Figure 2.**
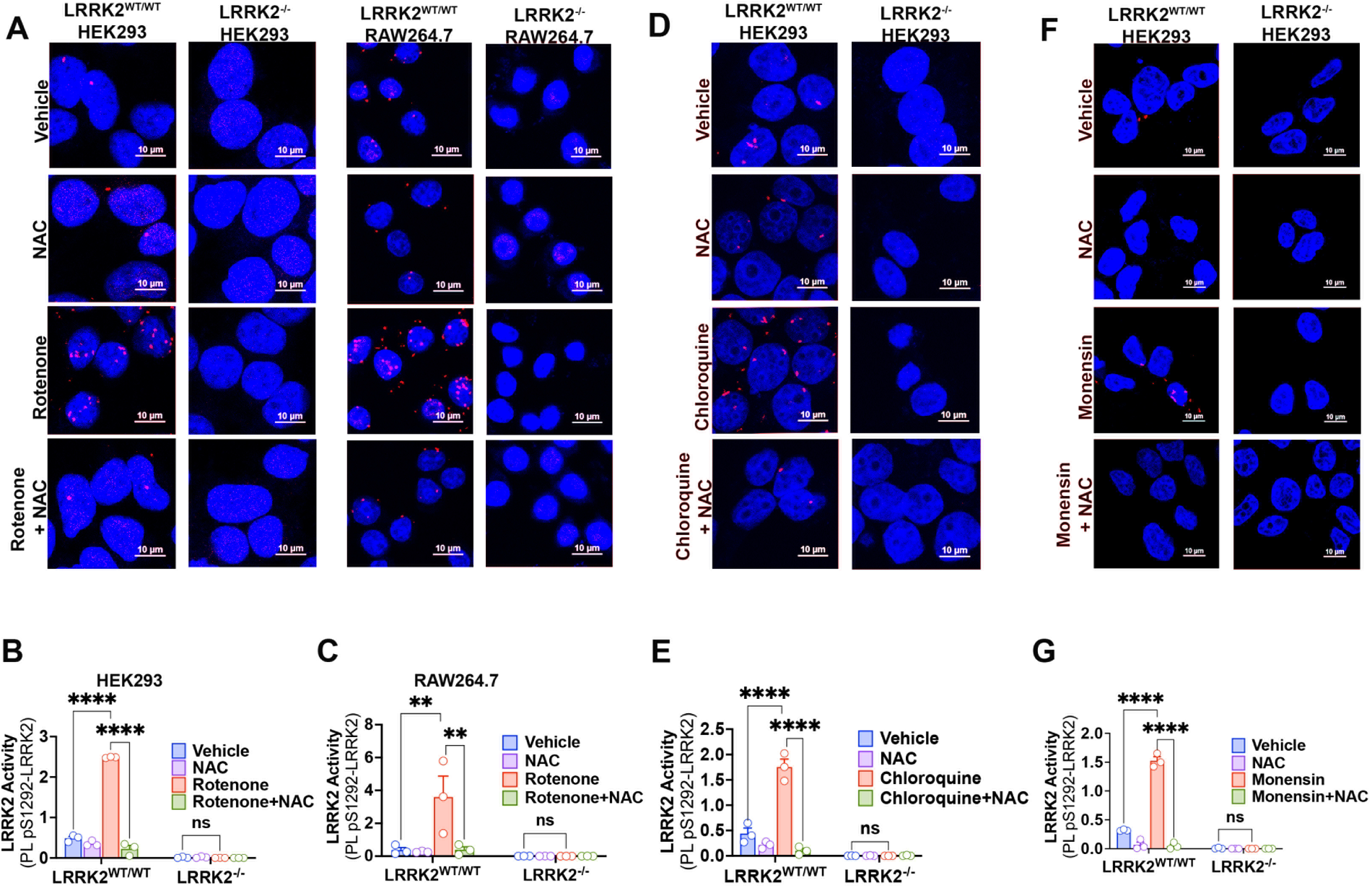
Stimulated LRRK2 activity is ROS dependent. (**A**) Rotenone (50nM) induced LRRK2 kinase activity was assessed by PL pS1292-LRRK2 in LRRK2^WT/WT^ and LRRK2^-/-^ HEK293 cells and RAW264.7 macrophages. Rotenone treatment led to an increase in PL pS1292-LRRK2 (red dots) in both LRRK2^WT/WT^ HEK293 and RAW264.7 macrophages compared to the vehicle. NAC co-treatment prevented rotenone-induced PL pS1292-LRRK2 signal and no signal was detected in LRRK2^-/-^ cells. (**B**) Quantification of LRRK2 activity by PL pS1292–LRRK2 Objects/DAPI in HEK293 cells. (**C**) Quantification of PL pS1292–LRRK2 Objects/DAPI in RAW264.7 macrophages. (**D**) Chloroquine (100μM) and (**F**) Monensin (10μM) elicited an increase in PL pS1292-LRRK2 signal in LRRK2^WT/WT^ HEK293 cells. NAC co-treatment prevented both (**D**) chloroquine and (**F**) monensin-induced PL pS1292-LRRK2 signal. No signal was observed in LRRK2^-/-^ cells. Quantification of LRRK2 activity by PL pS1292–LRRK2 in the presence of (**E**) chloroquine or (**G**) monensin. For each graph **(B, C, E, G)**, each symbol represents the average number of PL pS1292-LRRK2 objects/DAPI obtained from 100-150 cells per treatment group for each of three independent experiments. Statistical testing by two-way ANOVA with post-hoc Tukey correction. **denotes p<0.005; ****denotes p<0.0001; ns denotes not significant.

Given the high expression of LRRK2 in immune cells and its role in inflammatory responses in PD, we assessed whether rotenone-induced kinase activation was also redox-dependent in RAW264.7 macrophages. Consistent with the results in HEK293 cells, rotenone increased endogenous LRRK2 activity as measured by PL pS1292-LRRK2 in WT RAW264.7 macrophages (***Fig. 2A, C).*** Co-treatment with NAC prevented the rotenone-induced increase in PL pS1292– LRRK2 signal (p<0.005; two-way ANOVA with Tukey correction) (***Fig. 2A, C)***. Similar results were obtained using the orthogonal assays of pT73-Rab10 detection by western blot (p<0.05; two-way ANOVA with Tukey correction) (***fig. S1A, C)*** and PL pT73(Rab10)-Rab10 (p<0.0001; two-way ANOVA with Tukey correction) (***fig. S1D, F***). Treatment with NAC effectively blocked the rotenone-induced increase in pT73-Rab10 western blot signal (p<0.05; two-way ANOVA with Tukey correction) (***fig. S1A, C)*** and PL pT73(Rab10)-Rab10 signal (p<0.0001; two-way ANOVA with Tukey correction) (***fig. S1D, F***). Thus, using three separate assays (PL pS1292-LRRK2, pT73-Rab10 western blot, and PL pT73(Rab10)-Rab10) and two different cell lines (HEK293 and RAW264.7 macrophages), we found that rotenone stimulates endogenous LRRK2 kinase activity in a redox-dependent manner.

As noted, disruption of endolysosomal function or trafficking using chloroquine or monensin activates WT LRRK2 kinase (*21, 22, 35*). However, the mechanisms by which these disturbances in the endolysosomal system regulate LRRK2 kinase activity remain unclear. Given our findings with rotenone, we examined whether chloroquine or monensin activates LRRK2 kinase in a redox-dependent fashion. First, we assessed if chloroquine or monensin causes ROS production or oxidative damage. Using parameters known to elicit endogenous LRRK2 activity (chloroquine: 100 μM for 3 hours (*35*); monensin: 10 μM for 4 hours (*22*)) we measured cytoplasmic superoxide production using dihydroethidium (DHE). Under these conditions, both chloroquine and monensin elicited a significant increase in DHE signal compared to vehicle (p<0.005; one-way ANOVA with Tukey correction) (***fig. S3A, B***) in WT HEK293 cells. Thus, these stressors cause ROS production at concentrations and time points associated with elevated LRRK2 kinase activity.

As an orthogonal approach to assess if chloroquine and monensin lead to oxidative damage, we examined if they lead to the accumulation of the omega-6 PUFA-derived lipid hydroperoxidation end-product, 4-HNE. Consistent with the ROS production it induced, chloroquine caused an increase in 4-HNE at 1 hour (p<0.05; two-way ANOVA with Tukey correction) and 3 hours (p<0.001; two-way ANOVA with Tukey correction), which was prevented by co-treatment with NAC (***fig. S3E, F)***. Similarly, monensin caused an increase in 4-HNE at 4 hours (p<0.001; two-way ANOVA with Tukey correction), which was prevented by co-treatment with NAC (***fig. S3G, H)***. Given that these LRRK2 kinase activators cause ROS production and lipid peroxidation, both of which are prevented by NAC, we investigated whether NAC could prevent chloroquine- or monensin-induced LRRK2 kinase activity. Indeed, both chloroquine (***Fig. 2D, E***) and monensin (***Fig. 2F, G)*** treatment of WT HEK293 cells led to an increase in PL pS1292–LRRK2 signal (p<0.0001; two-way ANOVA with Tukey correction) that was prevented by NAC co-treatment. Together, these results indicate that known LRRK2 kinase activators, which result in either mitochondrial dysfunction (rotenone), lysosomal dyshomeostasis (chloroquine), or vesicular trafficking deficits (monensin), may activate LRRK2 through a common redox mechanism.

### Kinase activation loop cysteines 2024 and 2025 are important for redox-dependent activation of LRRK2

The dynamics of the kinase activation loop plays a critical role in driving LRRK2 kinase activity (*12, 13*). The activation loop contains two cysteine residues, Cys2024 and Cys2025 (*37*), that are highly conserved across many species (*38*) but not present in LRRK1 (*39*), suggesting a specific role in LRRK2 function. Cysteine residues on proteins can undergo modifications affecting function (*25–28*). We hypothesized that Cys2024 and Cys2025 are important for sensing the redox state of the local subcellular environment (*12*) and thereby for regulating LRRK2 kinase activation. Using CRISPR/Cas9, we generated HEK293 cell lines with each of these cysteines mutated individually to an alanine (LRRK2^C2024A^ and LRRK2^C2025A^) or together in a double mutant (LRRK2^C2024A+C2025A^). To ensure that we did not create “kinase-dead” mutants, baseline LRRK2 kinase activity was assessed using two assays: PL pS1292–LRRK2 and PL pT73(Rab10)-Rab10 (***fig S4A-D***). Under basal conditions, LRRK2^C2024A^, LRRK2^C2025A^, and LRRK2^C2024A+C2025A^ cells had PL pS1292-LRRK2 and PL pT73(Rab10)-Rab10 signals equivalent to LRRK2^WT/WT^ (***fig S4A-D***). Importantly, PF360 treatment decreased basal PL pS1292-LRRK2 and PL pT73(Rab10)-Rab10 signals (p<0.0001; two-way ANOVA with Tukey correction) (***fig S4A-D***) in all cell lines.

Having demonstrated that that baseline kinase activity of endogenous LRRK2 was not reduced in any of the cysteine mutant cell lines, we examined their responses to redox challenges. We found that while H_2_O_2_ exposure elevated PL pS1292–LRRK2 signal in WT HEK293 cells (p<0.0001; two-way ANOVA with Tukey correction), there was no increase in PL pS1292–LRRK2 signal in any of the cysteine mutant cell lines (***Fig. 3A, B)***. Next we explored if rotenone, chloroquine, or monensin were able to enhance kinase activity in LRRK2^C2024A^, LRRK2^C2025A^, and LRRK2^C20284A+C2025A^ HEK293 cells. In WT HEK293 cells, rotenone, chloroquine, and monensin treatment each led to an increase in PL pS1292–LRRK2 signal compared to vehicle (p<0.0001; two-way ANOVA with Tukey correction) (***Fig. 3C-H*).** Strikingly, in LRRK2^C2024A^, LRRK2^C2025A^, and LRRK2^C2024A+C2025A^ cells there was no increase in PL pS1292–LRRK2 signal in response to rotenone, chloroquine, or monensin (***Fig. 3C-H*)**.

**Figure 3.**
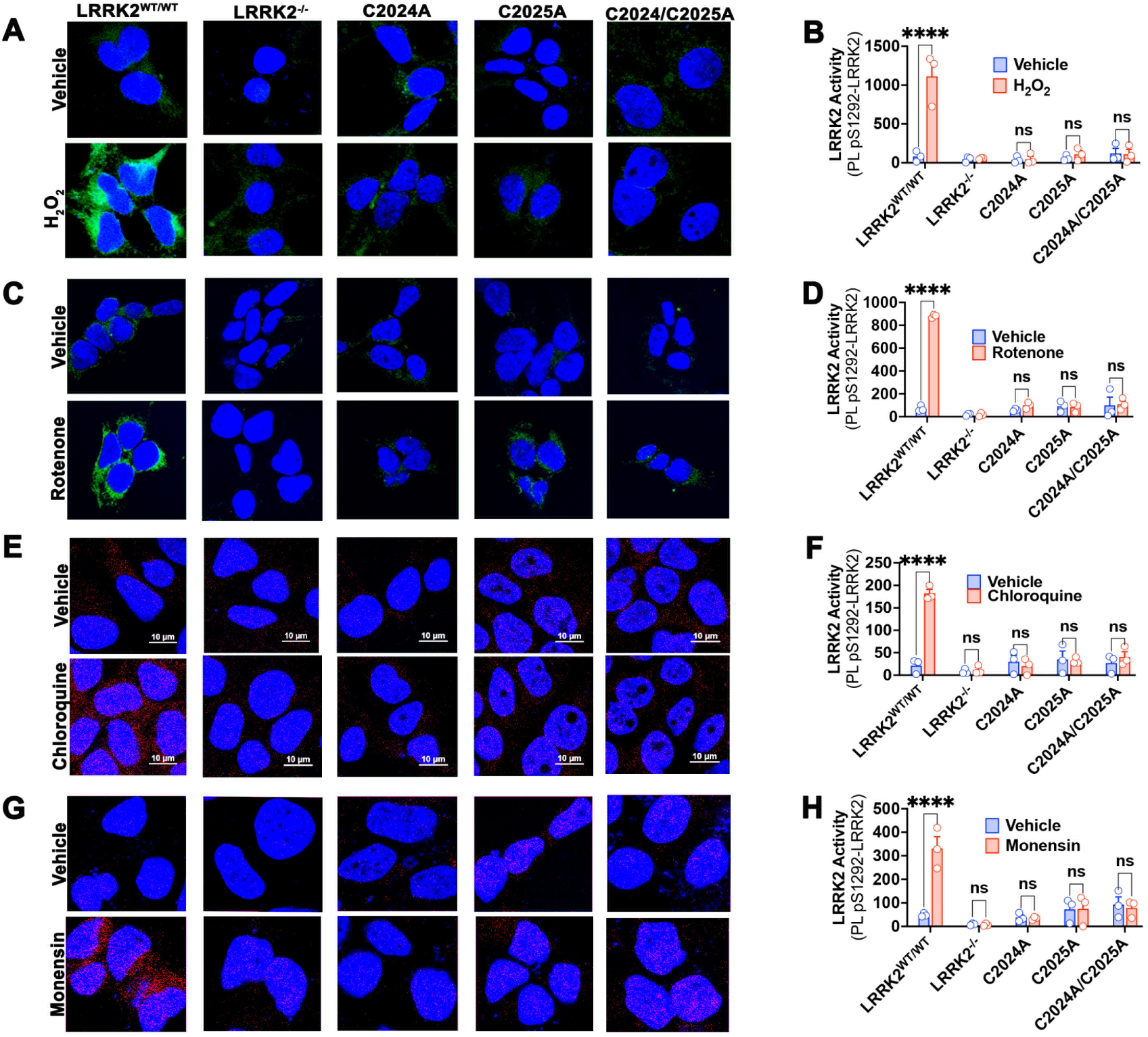
Cys2024 and Cys2025 are important for stimulating LRRK2 kinase activity. Endogenous LRRK2 kinase activity was assessed by PL pS1292-LRRK2 in LRRK2^WT/WT^, LRRK2^C2024A^, LRRK2^C2025A^, and LRRK2^C2024A+C2025A^ (denoted as C2024A, C2025A, C2024A/C2025A respectively) HEK293 cells. (**A**) H_2_O_2_ (5μM) (**C**) rotenone (50nM) (**E**) chloroquine (100μM) and (**G)** monensin (10μM) elicited an increase in PL pS1292-LRRK2 signal in LRRK2^WT/WT^, which was prevented in LRRK2^C2024A^, LRRK2^C2025A^, and LRRK2^C2024A+C2025A^ HEK293 cells. Note that the PL signal is green for H_2_O_2_ and rotenone and red for chloroquine and monensin. No signal was observed in LRRK2^-/-^ cells (**A, C, E, G**). Quantification of LRRK2 activity by PL pS1292–LRRK2 fluorescence signal in presence of (**B**) H_2_O_2_, (**D**) rotenone, (**F**) chloroquine, and (**H**) monensin. For all graphs **(B, D, F, H)**, each symbol represents the average PL pS1292–LRRK2 fluorescence intensity obtained from 100-150 cells per treatment group for each of three independent experiments. Statistical testing by two-way ANOVA with Tukey correction; ****denotes p<0.0001; ns denotes not significant.

It was previously shown that endogenous LRRK2 activity regulates rotenone-induced ROS production (*40, 41*). To assess whether mutating these cysteine residues (thereby blocking the ability of rotenone to stimulate LRRK2 kinase activity) had downstream effects, we measured rotenone-induced ROS production in each of the cell lines (***fig S5***). Rotenone treatment led to an increase in DHE signal in WT HEK293 cells (p<0.0001; two-way ANOVA with Tukey correction) but not in LRRK2^C2024A^, LRRK2^C2025A^ or LRRK2^C2024A+C2025A^ cells (***fig S5***). Together, these results indicate that Cys2024 and Cys2025 are important residues that regulate kinase activation in a redox-responsive fashion.

### 4-HNE is a critical signaling mediator of LRRK2 kinase activity

LRRK2 kinase activity is provoked by membrane-damaging agents (*19, 20*) that cause lipid hydroperoxidation and accumulation of 4-HNE, a reactive aldehyde known to activate several kinases, including ERK, JNK, Src, and p38 MAPK (*25–28*). We examined whether exogenous 4-HNE could interact with and activate endogenous WT LRRK2 kinase activity. For this purpose, we devised a new PL assay, PL LRRK2-4-HNE, using antibodies that recognize 4-HNE and total LRRK2 to detect the formation of 4-HNE-LRRK2 adducts. When WT HEK293 cells were treated with a pathophysiological concentration of 4-HNE (100 μM for 1 hour), there was an increase in PL LRRK2–4-HNE signal, which was prevented by pre-treatment with NAC (p<0.0001; two-way ANOVA with Tukey correction) (***fig. S6***). This correlated with increased endogenous WT LRRK2 kinase activity as measured by PL pS1292-LRRK2 (p<0.005; two-way ANOVA with Tukey correction) (***fig. S7A, B***), and PL pT73(Rab10)-Rab10 (p<0.005; two-way ANOVA with Tukey correction (***fig. S7C, D***). NAC pre-treatment prevented the 4-HNE-induced increase in both PL pS1292-LRRK2 (***fig. S7A, B***), and PL pT73(Rab10)-Rab10 (***fig S7C, D***). These results demonstrate that exogenous 4-HNE can form adducts with and activate endogenous WT LRRK2.

To further examine the relationship between LRRK2–4-HNE adducts and LRRK2 kinase activity, WT HEK293 cells were treated with varying concentrations of 4-HNE (10 – 100 µM for 1 hour) and LRRK2–4-HNE adducts and LRRK2 activity were measured by PL. Increasing concentrations of 4-HNE caused parallel, dose-dependent increases in PL LRRK2–4-HNE and PL pS1292-LRRK2 signals (***Fig. 4A-D***) in WT HEK293 cells. In fact, within this concentration range, there was a highly significant linear relationship between the degree of LRRK2–4-HNE adduct formation and LRRK2 kinase activity (p<0.005; R^2^=0.99) (***fig. S8***). Given that 4-HNE has a high affinity for forming irreversible Michael adducts with cysteine residues, we also investigated whether the cysteine mutants influence LRRK2-4-HNE adduct formation and 4-HNE-induced kinase activity. Surprisingly, in LRRK2^C2024A+C2025A^ HEK293 cells, 4-HNE treatment did not increase PL LRRK2-4-HNE or PL pS1292-LRRK2 signals (***Fig 4A-D***). This suggests that, within this concentration range, 4-HNE binds preferentially to the Cys2024 and Cys2025 residues of LRRK2 in a cellular context.

**Figure 4.**
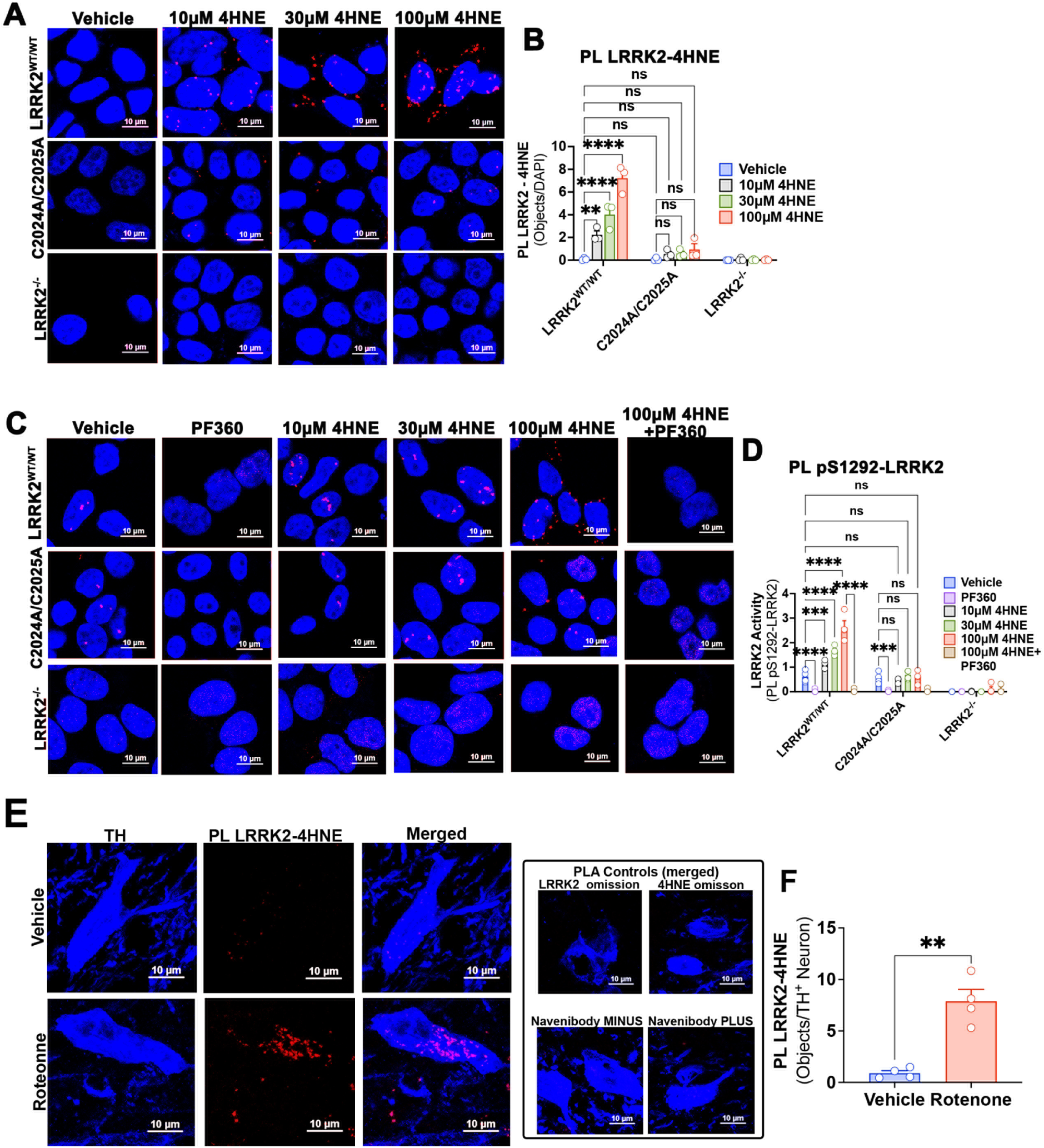
The lipid hydroperoxidation product 4-HNE interacts with LRRK2 and induces kinase activation. (**A**) PL between LRRK2 and 4-HNE (PL LRRK2–4-HNE) was used as an index of LRRK2-4-HNE adduct formation in LRRK2^WT/WT^, C2024A/C2025A, and LRRK2^-/-^ HEK293 cells. Exogenous 4-HNE treatment (10μM, 30μM, 100μM) led to a dose-dependent increase in PL LRRK2–4-HNE signal (red dots) in LRRK2^WT/WT^ cells. 4-HNE treatment did not increase PL LRRK2–4-HNE signal in C2024A/C2025A and LRRK2^-/-^ HEK293 cells. (**B**) Quantification of PL LRRK2-4-HNE. (**C**) 4-HNE-induced LRRK2 kinase activity was assessed by PL pS1292-LRRK2. Baseline kinase activity in LRRK2^WT/WT^ and C2024A/C2025A cells was inhibited by the LRRK2 kinase inhibitor PF360 (1μM), and 4-HNE treatment dose-dependently increased PL pS1292-LRRK2 signal (red dots) in LRRK2^WT/WT^ cells, but not in C2024A/C2025A cells. (**D**) Quantification of LRRK2 activity by PL pS1292–LRRK2 Objects/DAPI in HEK293 cells. For graphs **(B** and **D)**, each symbol represents the average signal from 100-150 cells per treatment group for each of three independent experiments. Statistical testing by two-way ANOVA with post-hoc Tukey correction. (**E**) LRRK2–4-HNE interaction was assessed via PL LRRK2–4-HNE (red dots) in TH^+^ dopaminergic neurons (blue) in in the substantia nigra of rats treated with vehicle or rotenone. Rotenone treatment caused an increase in PL LRRK2–4-HNE in TH^+^ dopaminergic neurons compared to vehicle treated rats. (**F**) Quantification of the PL LRRK2–4-HNE signal in dopamine neurons. Each symbol represents the average number of PL LRRK2–4-HNE objects/TH^+^ neuron from an individual animal with two sections of brain tissue stained per animal; N=4 animals per treatment group. Statistical testing by two-tailed unpaired *t*-test with Welch’s correction. **denotes p<0.01; ***denotes p<0.0005; ****denotes p<0.0001; ns denotes not significant.

To explore if LRRK2–4-HNE adduct formation occurs with endogenous generation of 4-HNE in response to stimuli that activate LRRK2 kinase activity, we examined the effects of rotenone, chloroquine, and monensin. Compared to vehicle, rotenone elicited an increase in PL LRRK2–4-HNE signal in LRRK2^WT/WT^ HEK293 (p<0.0001; two-way ANOVA with Tukey correction) and RAW267.4 macrophages (p<0.0001; two-way ANOVA with Tukey correction) (***fig. S9)***. Co-treatment with α-tocopherol, a compound that can protect against lipid hydroperoxidation (*42, 43*), prevented the rotenone-induced increase in PL LRRK2–4-HNE signal and no signal was observed in LRRK2^-/-^ cells (***fig. S9)***. Similar to rotenone, both chloroquine and monensin treatment led to an increase in PL LRRK2–4-HNE signal compared to vehicle (p<0.0001; two-way ANOVA with Tukey correction) (***fig. S10***). Co-treatment with α-tocopherol prevented the chloroquine- and monensin-induced increase in PL LRRK2–4-HNE and no signal was observed in LRRK2^-/-^ cells (***fig. S10***).

We previously showed there is 4-HNE accumulation and elevated LRRK2 kinase activity in dopaminergic neurons of the substantia nigra (SN) in the rotenone rat model of PD (*9, 44, 45*). Therefore, we investigated whether LRRK2–4-HNE adduct formation also occurred in this model. As described (*45*), middle-aged rats were treated with rotenone for 7-10 days until they reached behavioral endpoint, at which time they were euthanized for analysis. Rotenone treatment led to a marked increase in PL LRRK2–4-HNE signal in SN dopamine neurons compared to vehicle ***(Fig. 4E, G)***) (p<0.01; two tailed unpaired *t*-test with Welch’s correction), thereby providing *in vivo* relevance of LRRK2–4-HNE adduct formation. Together, these data demonstrate that the lipid hydroperoxidation end-product, 4-HNE, may be a key signaling mediator in LRRK2 kinase activation that links mitochondrial and endolysosomal stress to LRRK2 kinase activation.

### 15-Lipoxygenase regulates LRRK2 kinase activity

Since 4-HNE forms adducts with LRRK2 and regulates its kinase activation, we explored the possible upstream sources of this reactive aldehyde. 4-HNE can be produced nonenzymatically when membrane lipids react directly with ROS; alternatively, it can be produced enzymatically as the end-product of 15-LO activity (*29, 30*). Our data demonstrate that α-tocopherol can prevent stimulus-induced PL LRRK2–4-HNE in an endogenous context, but it is unclear whether this is due to its ROS scavenging properties or its ability to inhibit 15-LO (*33*). We therefore used both genetic and pharmacological approaches to assess the role of 15-LO in producing the pool of 4-HNE that activates LRRK2. We utilized CRISPR/Cas9 gene-edited 15-LO knockout (15-LO^-/-^) HAP1 cells and a novel, potent, and selective small molecule 15-LO inhibitor, CU-12991, that has an IC_50_ of 2.5 nM for inhibition of rotenone-induced 4-HNE production (***fig. S11***). In WT cells, rotenone elicited an increase in cellular 4-HNE, which was prevented by co-treatment with 20nM CU-12991 (p < 0.0001; two-way ANOVA with Tukey correction) **(*fig. S12A, B)***. There was no rotenone-induced 4-HNE accumulation in 15-LO^-/-^ cells, and CU-12991 had no effect in these cells **(*fig. S12A, B)***. Thus, in the absence of 15-LO activity, there was no rotenone-induced 4-HNE production.

We next explored whether 15-LO-derived 4-HNE mediates rotenone-induced LRRK2 activity. First, we assessed LRRK2–4-HNE adduct formation via PL LRRK2–4-HNE. Consistent with our previous observations (***Fig. 4 and Fig. S9),*** rotenone increased the PL LRRK2–4-HNE signal compared to the vehicle (p<0.0005; two-way ANOVA with Tukey correction), and this was prevented by co-treatment with α-tocopherol (***Fig. 5A, B***). Strikingly, there was no rotenone-induced PL LRRK2–4-HNE signal in 15-LO^-/-^ cells (***Fig. 5A, B***). Next, we assessed kinase activation by both autophosphorylation (***Fig. 5C, D***) and substrate phosphorylation (***fig. S12C, D).*** In WT cells, rotenone increased PL pS1292-LRRK2 signal (p<0.0001; two-way ANOVA with Tukey correction) (***Fig. 5C, D***) and PL pT73(Rab10)-Rab10 signal (p<0.0001; two-way ANOVA with Tukey correction) (***fig. S12C, D)***. Interestingly, 15-LO inhibition by CU-12991 mitigated the rotenone-induced increase in PL pS1292-LRRK2 (***Fig. 5C, D***) and PL pT73(Rab10)-Rab10 (***fig. S12C, D)***, but did not reduce LRRK2 kinase activity below basal levels (***Fig. 5C, D***). In contrast, PF360 treatment completely abolished both basal and rotenone-induced kinase activity (PL pS1292-LRRK2, ***Fig. 5C, D***) (PL pT73(Rab10)-Rab10, ***fig. S12C, D***).

**Figure 5.**
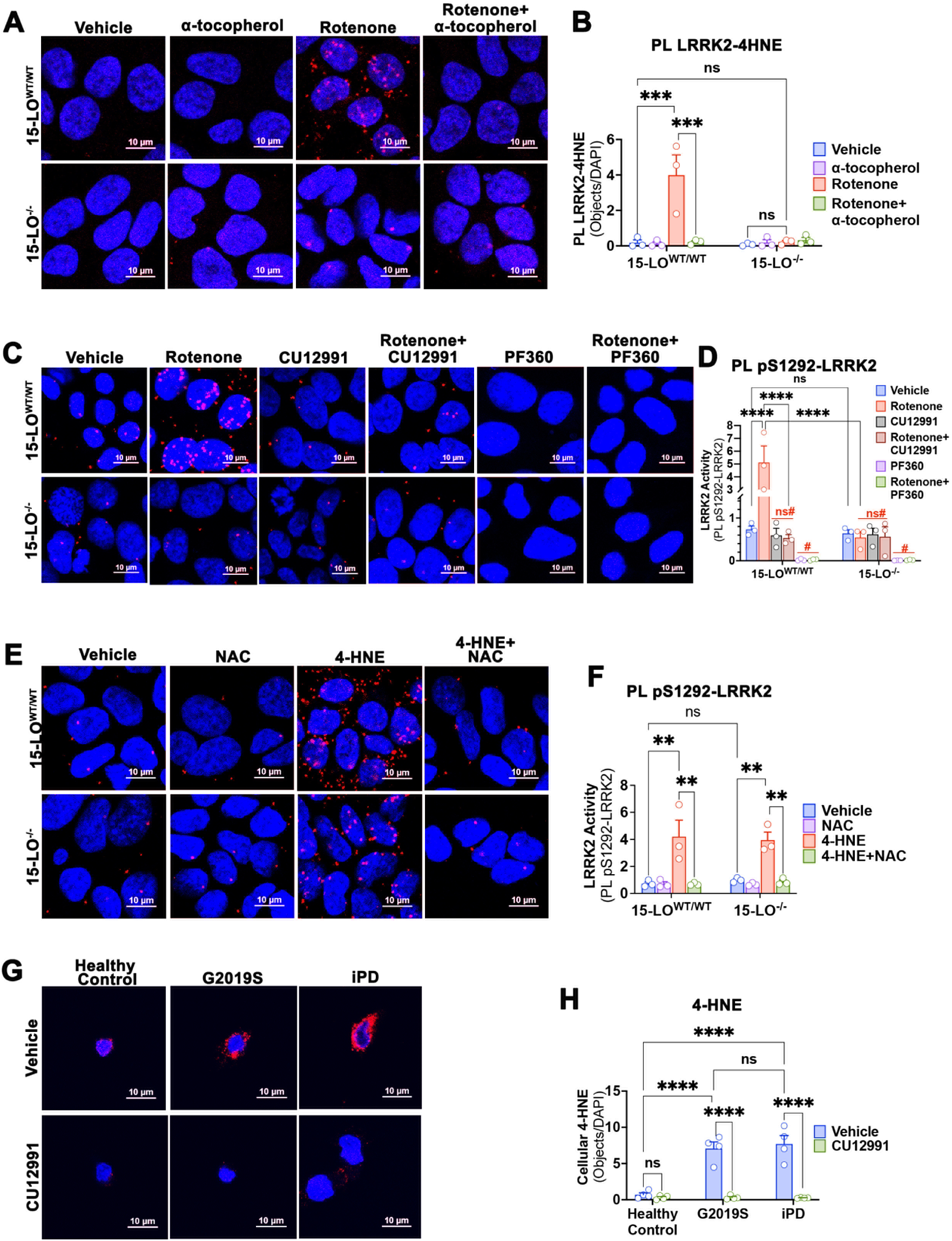
15-LO-derived 4-HNE is a key mediator of LRRK2 hyperactivation. (**A**) Rotenone-induced endogenous LRRK2–4-HNE adduct formation was assessed by PL LRRK2-4-HNE in 15-LO^WT/WT^ and 15-LO^-/-^ HAP1 cells. Rotenone treatment promoted an increase in PL LRRK2-4-HNE signal that was prevented by α-tocopherol in 15-LO^WT/WT^, and not observed in 15-LO^-/-^ cells. (**B**) Quantification of PL LRRK2-4-HNE. (**C**) Rotenone-induced LRRK2 kinase activity was assessed by PL pS1292-LRRK2. Rotenone increased PL pS1292-LRRK2, an effect that was that was mitigated by cotreatment with the 15-LO inhibitor, CU-12991, and completely prevented by PF360. 15-LO^-/-^ cells had a normal basal PL pS1292 – LRRK2 signal, which was not further increased by rotenone or affected by CU-12991. (**D**) Quantification of LRRK2 activity by PL pS1292-LRRK2. (**E**) 4-HNE (100μM) treatment led to an increase in PL pS1292-LRRK2 signal in both 15-LO^WT/WT^ and 15-LO^-/-^ cells. This was prevented by pretreatment with 250μM NAC. (**F**) Quantification of LRRK2 activity by PL pS1292-LRRK2. For graphs **(B, D,** and **F)**, each symbol represents the average signal from 100-150 cells per treatment group for each of three independent experiments. Statistical testing by two-way ANOVA with post-hoc Tukey correction. (**G**) Cellular 4-HNE signal in LCLs derived from healthy control subjects, G2019S mutation carriers, and iPD patients. The increased 4-HNE signal in G2019S and iPD LCLs was reduced to healthy control levels by CU-12991. (**H**) Quantification of 4-HNE. Each symbol represents an individual donor line. 4-HNE signal was taken from 90-120 cells/line; 4 donor lines per group. Statistical testing by two-way ANOVA with post-hoc Tukey correction. **denotes p<0.005; *** denotes p<0.0005; ****denotes p<0.0001; ns denotes not significant. For panel D, red symbols indicate additional statistical testing by one-way ANOVA, examining each genotype (15-LO^WT/WT^ and 15-LO^-/-^) separately. ns# denotes not significant compared to vehicle-treated cells; # denotes p< 0.05 compared to vehicle-treated or CU-12991-treated cells.

Consistent with our results using CU-12991, in 15-LO^-/-^ cells there was no rotenone-induced PL pS1292-LRRK2 (***Fig. 5C, D***) or PL pT73(Rab10)-Rab10 (***fig. S12C, D***), although basal kinase activity was preserved. Importantly, CU-12991 did not impact LRRK2 activity in 15-LO^-/-^ cells ***(Fig 5C, D*** and ***fig. S12C, D***). Furthermore, to ensure CU-12991 was not directly interfering with 4-HNE, we treated cells with exogenous 4-HNE and found that induced kinase activity was unaffected by CU-12991 (***fig. S13)*.**

To ensure that the stimulatory effect of 15-LO on LRRK2 kinase activity was due to its production of 4-HNE, we treated cells with exogenous 4-HNE (***Fig. 5E, F***). In both WT and 15-LO^-/-^ cells, 4-HNE elicited an increase in PL pS1292-LRRK2 signal (p<0.005; two-way ANOVA with Tukey correction), which was prevented by treatment with NAC (***Fig 5E, F***). Thus, the effects of pharmacological inhibition or genetic deletion of 15-LO on LRRK2 kinase activity can be overcome by exogenous 4-HNE. Collectively, these results suggest that 4-HNE generated by 15-LO forms adducts with LRRK2, leading to its hyperactivation. Therefore, LRRK2 kinase activation is specifically regulated by the 4-HNE produced by 15-LO as an end-product of lipid hydroperoxidation.

Accumulation of 4-HNE in the SN of PD post-mortem human brain tissue has been recognized for 30 years (*46*); however, the functional role of 15-LO-derived 4-HNE in PD has yet to be explored. Therefore, we assessed 4-HNE levels in patient-derived lymphoblastoid cell lines (LCLs) derived from healthy control subjects, G2019S-LRRK2 PD patients, and iPD patients. Relative to healthy controls, basal 4-HNE was elevated in LCLs derived from subjects with iPD as well as subjects harboring a G2019S mutation (p < 0.0001 for both iPD and G2019S compared to healthy controls; two-way ANOVA with Tukey correction) ***(Fig. 5H)***. The elevated 4-HNE signal in the iPD and G2019S PD subjects was reduced to healthy control levels by treatment with CU-12991 (p < 0.0001 for both iPD and G2019S; two-way ANOVA with Tukey correction two-way ANOVA with Tukey correction) ***(Fig. 5H)***. This suggests that 15-LO is responsible for the pathological accumulation of 4-HNE observed in G2019S and iPD patient samples.

Next we investigated whether 15-LO-derived 4-HNE was responsible for LRRK2–4-HNE adduct formation elicited by other known LRRK2 activators: chloroquine, monensin, or rotenone. Consistent with our results in HEK cells *(**fig. S9, fig. S10)***, in WT RAW264.7 macrophages chloroquine, monensin, and rotenone each elicited an increase in PL LRRK2-4-HNE signal (***Fig. 6A, B***). As observed in 15-LO^-/-^ HAP1 cells (***Fig. 5A, B)***, 15-LO inhibition with CU-12991 prevented the formation of rotenone-induced LRRK2–4-HNE adducts (***Fig. 6A, B)*** in WT RAW264.7 macrophages. Similarly, CU-12991 co-treatment prevented chloroquine- and monensin-induced PL LRRK2-4-HNE signal in WT RAW264.7 macrophages (***Fig. 6A, B)***. Under the same conditions, chloroquine, monensin, and rotenone each significantly increased PL pS1292-LRRK2 signal in WT RAW264.7 macrophages (***Fig. 6C, D***). Strikingly, CU-12991 co-treatment completely prevented rotenone-, chloroquine-, and monensin-induced PL pS1292-LRRK2 signal (***Fig. 6C, D).*** As an orthogonal approach to measure LRRK2 kinase activity, we assessed substrate phosphorylation via pT73-Rab10 western blot. Chloroquine, monensin, and rotenone each led to an increase in pT73-Rab10, which was prevented by co-treatment with CU-12991 (***fig. S14***). Thus, using genetic and pharmacological approaches and three independent LRRK2 activity assays (PL pS1292–LRRK2, pThr-73-Rab10 western blot, and PL pT73(Rab10)-Rab10), our results indicate that endogenous stimulated LRRK2 activity is regulated by 15-LO.

**Figure 6.**
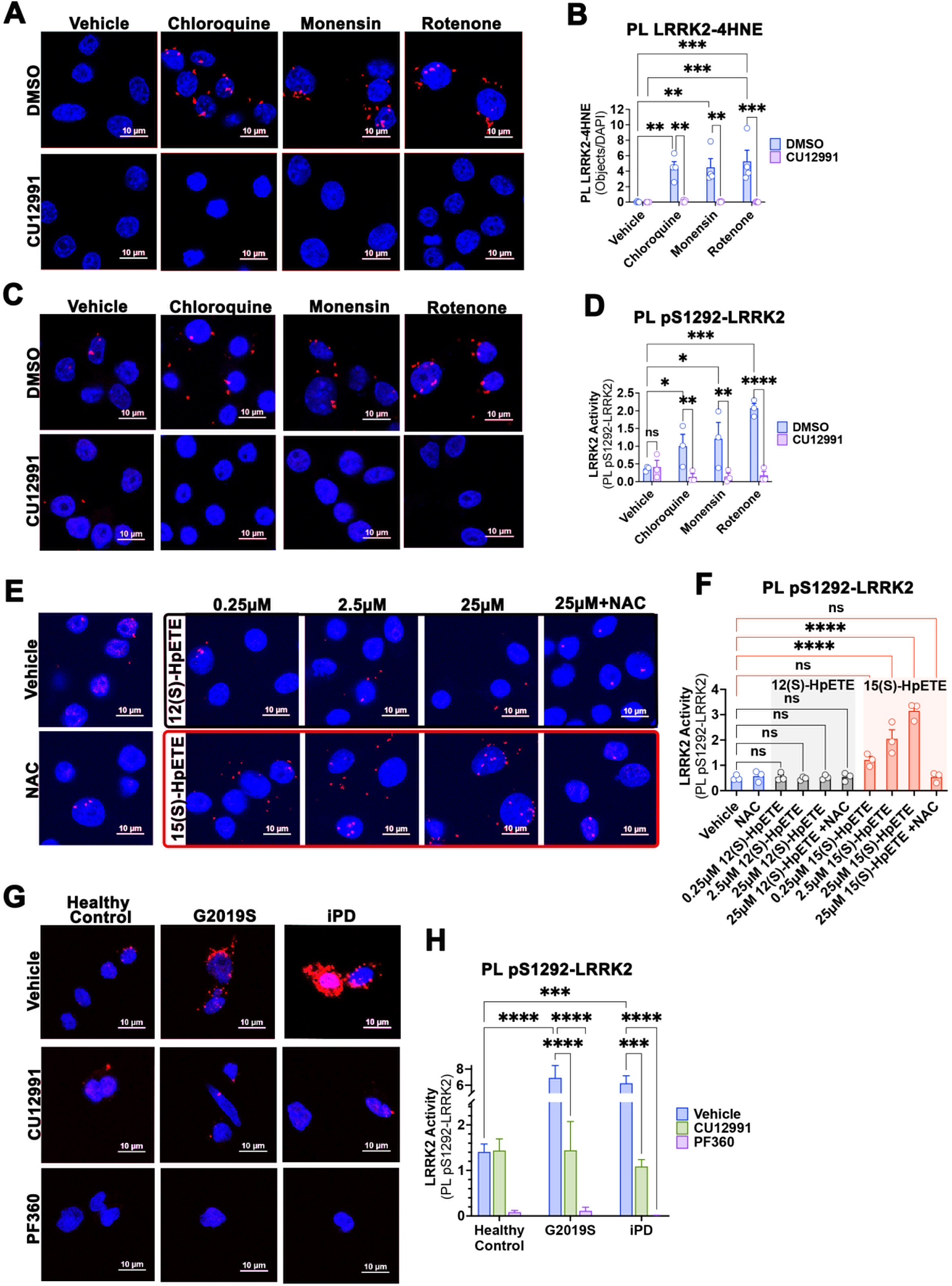
15-LO activity regulates stimulated LRRK2 kinase activity via 4-HNE. (**A**) LRRK2-4-HNE adduct formation was assessed by PL LRRK2-4-HNE in LRRK2^WT/WT^ RAW264.7 macrophages. Chloroquine, monensin, and rotenone led to an increase PL LRRK2-4-HNE signal (red dots). The increase in PL LRRK2-4-HNE signal was prevented by cotreatment with CU-12991. (**B**) Quantification of PL LRRK2-4-HNE. (**C**) Chloroquine, monensin, and rotenone-induced kinase activity was assessed by PL pS1292-LRRK2 in LRRK2^WT/WT^ RAW264.7 macrophages. Chloroquine, monensin and rotenone caused increased PL pS1292-LRRK2 signal (red dots) which was prevented by cotreatment with the 15-LO inhibitor, CU-12991. (**D**) Quantification of LRRK2 activity by PL pS1292-LRRK2. (**E**) LRRK2^WT/WT^ RAW264.7 macrophages were treated with either 12(S)-HpETE (top panel, black outline) or 15(S)-HpETE (bottom panel, red outline) and kinase activity was measured by PL pS1292-LRRK2. 12(S)-HpETE did not increase PL pS1292-LRRK2 signal above baseline (red dots). 2.5μM and 25μM 15(S)-HpETE led to an increase in PL pS1292-LRRK2 signal relative to vehicle. Cotreatment with NAC prevented the increase in PL pS1292-LRRK2 signal elicited by 25μM 15(S)-HpETE. (**F**) Quantification of LRRK2 activity by PL pS1292–LRRK2. For each graph (**B**, **D**, and **F**), each symbol represents the average number of PL LRRK2-4-HNE Objects/DAPI or PL pS1292-LRRK2 Objects/DAPI obtained from 100-150 cells per treatment group for each of 3-4 independent experiments. Statistical testing by two-way ANOVA with post-hoc Tukey correction. (**G**) LRRK2 activity measured by PL pS1292-LRRK2 in LCLs derived from healthy control subjects, G2019S mutation carriers, and iPD patients. The increased PL pS1292-LRRK2 signal in G2019S and iPD LCLs was reduced to healthy control basal levels by CU-12991 and completely blocked by PF360. (**H**) Quantification of LRRK2 activity by PL pS1292-LRRK2 Objects/DAPI was taken from 90-120 cells/line; 4 donor lines per group. Statistical testing by two-way ANOVA with post-hoc Tukey correction. * denotes p<0.05; **denotes p<0.005; *** denotes p≤0.0005; **** denotes p<0.0001; ns denotes not significant.

Although both 15-LO and 12-lipoxygenase (12-LO) can participate in lipid peroxidation, 15-LO activity is the predominant source of 4-HNE formation. The initial step in this enzymatic pathway (for both 15-LO and 12-LO) produces their respective hydroperoxy lipid metabolites: 15(S)-HpETE for 15-LO and 12(S)-HpETE for 12-LO (*29, 30*). A subsequent intramolecular rearrangement of 15(S)-HpETE, but not 12(S)-HpETE, produces 4-HNE (*47*). To further validate the 15-LO pathway and subsequent 4-HNE formation as a regulator of LRRK2 kinase activity, we treated WT RAW264.7 macrophages with either 15(S)-HpETE or 12(S)-HpETE and assessed endogenous LRRK2 kinase activation by PL pS1292-LRRK2. 12(S)-HpETE treatment (0.25 – 25 µM) did not increase PL pS1292–LRRK2 signal (***Fig. 6E, F***); however, 15(S)-HpETE dose-dependently increased PL pS1292–LRRK2 signal (p<0.0001; two-way ANOVA with Tukey correction) (***Fig. 6E, F*).** NAC pre-treatment prevented 15(S)-HpETE-induced PL pS1292– LRRK2 signal (***Fig. 6E, F*)**. Together, these results indicate that LRRK2 kinase activation was driven by 15(S)-HpETE rearrangement to 4-HNE.

To determine the PD relevance of 15-LO-regulated LRRK2 activity, we next investigated if LRRK2 activity was elevated in G2019S PD and iPD patient-derived LCLs. Relative to healthy controls, there was elevated PL pS1292-LRRK2 in both G2019S (p < 0.0001; two-way ANOVA with Tukey correction) and iPD LCLs (p < 0.005; two-way ANOVA with Tukey correction) that was completely abolished by PF360 (***Fig. 6G, H)***. Strikingly, CU-12991 reduced pathologically hyperactive LRRK2 kinase activity in both G2019S and iPD LCLs back to basal healthy control levels but did not completely inhibit LRRK2 activity (***Fig. 6G, H***). Collectively, these data provide strong evidence that 15-LO activity regulates pathologically elevated endogenous LRRK2 kinase activity through the enzymatic production of 4-HNE and its subsequent adduct formation with LRRK2.

The ability of 15-LO to hydroperoxidate membrane phospholipids depends on its association with the small scaffolding protein, phosphatidylethanolamine-binding protein (PEBP1) (*48*). Typically, PEBP1 is in complex with RAF1 kinase; however, upon phosphorylation, it is liberated and interacts with new proteins (*49*), including 15-LO. In the unbound state, 15-LO prefers as substrates free fatty acids, not those incorporated into membranes. However, when complexed with PEBP1, its substrate preference switches to omega-6 PUFAs incorporated into membrane phosphatidylethanolamines (*48, 50*), and omega-6 PUFA hydroperoxidation results in the production of 4-HNE. We therefore explored the interaction between PEBP1 and 15-LO in a system (rotenone-treated RAW264.7 macrophages) where endogenous 4-HNE production is stimulated with resultant LRRK2 kinase activation. Quantitative confocal immunofluorescence staining revealed that rotenone treatment markedly increased the colocalization of the PEBP1 and 15-LO (p < 0.005; two tailed unpaired *t*-test with Welch’s correction) (***Fig. 7A, B***). Using a complementary approach, we developed a PL assay that detects the interaction between PEBP1 and 15-LO (PL PEBP1–15-LO). Similar to our colocalization results, rotenone-induced a strong PL PEBP1–15-LO signal compared to vehicle (***Fig. 7C, D)***.

**Figure 7.**
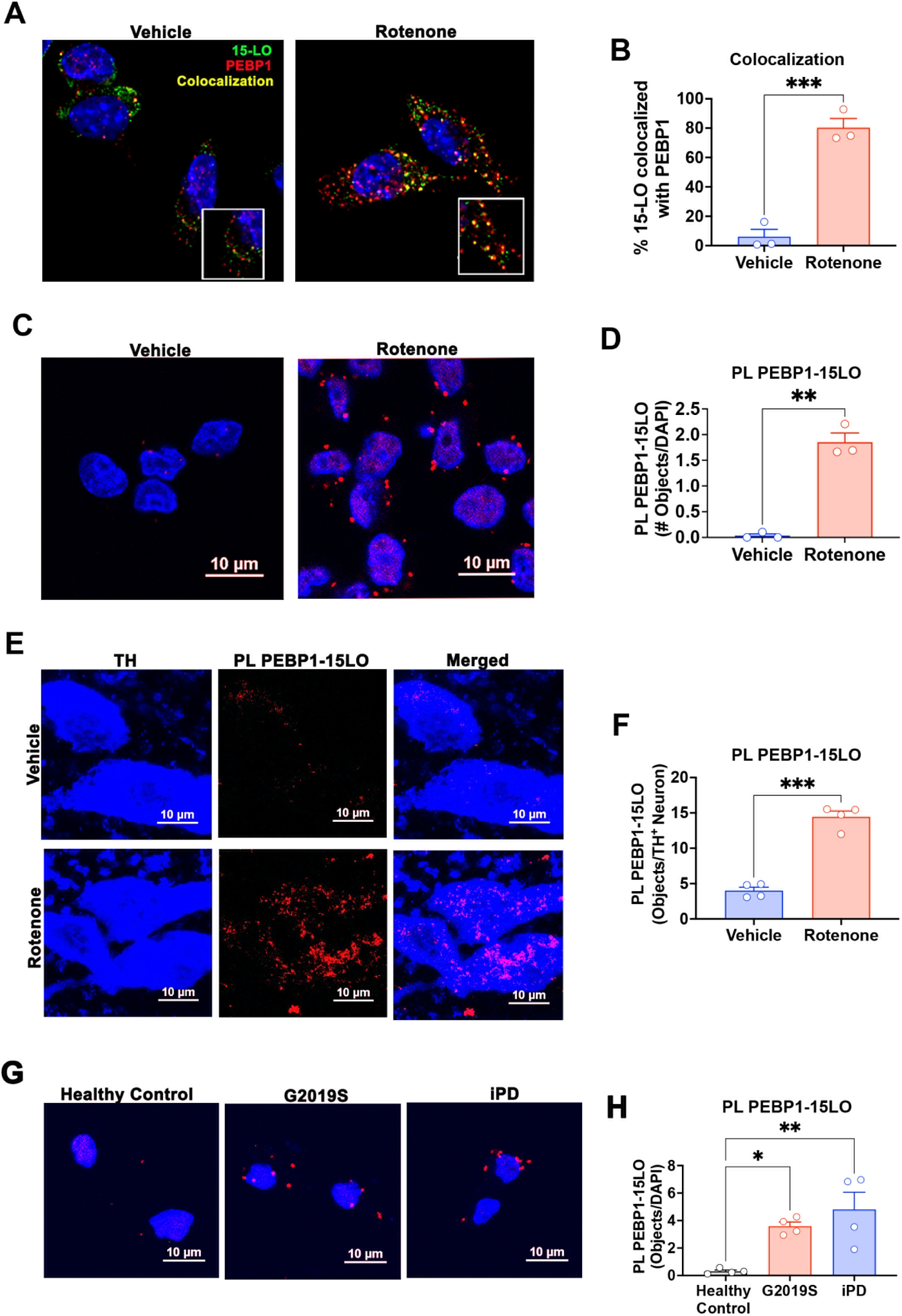
PEBP1 and 15-LO interaction is increased in the rotenone model and patient derived samples. (**A**) Immunofluorescent staining for 15-LO (green), PEBP1 (red), and their colocalization (yellow) in LRRK2^WT/WT^ RAW264.7 macrophages. Rotenone led to an increase in overlap between 15-LO and PEPB1 relative to vehicle. Inset is 200X zoom. (**B**) Quantification of the % overlap of 15-LO and PEBP1 objects/Total Number of 15-LO Objects. Each symbol represents the average % overlap of 15-LO and PEBP1 objects/Total Number of 15-LO Objects obtained from 100-150 cells per treatment group for each of three independent experiments. Statistical testing by two tailed unpaired *t*-test with Welch’s correction. *** denotes p<0.001 (**C**) PL between PEBP1 and 15-LO was used as an index for PEPB1–15-LO interaction in LRRK2^WT/WT^ RAW264.7 macrophages. Rotenone treatment led to an increase in PL PEBP1–5-LO signal (red dots) compared to vehicle. (**D**) Quantification of PL PEBP1–15-LO. Each symbol represents the average number of PL PEBP1–15-LO Objects/DAPI obtained from 100-150 cells per treatment group for each of three independent experiments. Statistical testing by two tailed unpaired *t*-test with Welch’s correction. ** denotes p<0.01 (**E**) PEBP1–15-LO interaction was assessed via PL PEBP1–15-LO (red dots) in TH^+^ dopaminergic neurons (blue) in the substantia nigra of rats treated with vehicle or rotenone. Rotenone treatment caused an increase in PL PEBP1–15-LO in TH^+^ dopaminergic neurons compared to vehicle-treated rats. (**F**) Quantification of the PL PEBP1–15-LO signal in dopamine neurons. Each symbol represents the average number of PL PEBP1–15-LO HNE objects/TH^+^ neuron from an individual animal with two sections of brain tissue stained per animal; N=4 animals per treatment group. Statistical testing by two-tailed unpaired *t*-test with Welch’s correction.***denotes p<0.0005. (**G**) PL PEBP1–15-LO was assessed in LCLs derived from healthy control subjects, G2019S mutation carriers, and iPD patients. There was an increase in PL PEBP1–15-LO signal in G2019S and iPD derived LCLs relative to healthy control. (**H**) Quantification of PL PEBP1–15-LO Objects/DAPI. Each symbol represents the average number of PL PEBP1–15-LO Objects/DAPI taken from 90-120 cells/line; 4 donor lines per group. Statistical testing by one-way ANOVA with post-hoc Tukey correction * denotes p<0.05; ** denotes p≤0.005.

We also examined if this occurred *in vivo* in the rotenone rat model of PD (***Fig. 7E, F***). Analogous to our findings in macrophages, there was an increase in PL PEBP1–15-LO signal in dopaminergic neurons in rotenone-treated rats compared to vehicle (p < 0.0005; two-tailed unpaired *t*-test with Welch’s correction) (***Fig. 7E, F***). To investigate if this was relevant in the human disease, we sought to determine if PL PEBP1-15-LO was increased in LCLs derived from G2019S PD and iPD patients compared to healthy controls. Indeed, there was an elevated PL PEBP1–15-LO signal in LCLs derived from G2019S (p < 0.05; one-way ANOVA with Tukey correction) and iPD (p=0.005; one-way ANOVA with Tukey correction) patients compared to healthy controls (***Fig. 7G, H***). Thus, using complementary approaches in multiple model systems and PD patient samples, the data demonstrate that under conditions generating endogenous 4-HNE and stimulating LRRK2 kinase activity, there is an enhanced association between 15-LO and PEBP1.

## DISCUSSION

Pathogenic mutations in *LRRK2* that cause elevated kinase activity are predominantly driven by altered kinetics, altered GTP hydrolysis, or enhanced substrate specificity (*8, 12, 13, 51*). Although each pathogenic mutation can increase LRRK2 kinase activity, they do so in distinct ways, highlighting the complexity of LRRK2 activation. When assessing kinase activity, cellular *in situ* assays have advantages over *in vitro* kinase assays, as the cellular context is important for LRRK2 activation (*6*). Upstream signaling events may be critical for understanding how and where endogenous WT LRRK2 activation occurs and what substrates are involved. In this work, we aimed to elucidate what regulates pathologically elevated LRRK2 kinase activity. To strengthen our conclusions, kinase activity was assessed using three complementary approaches: (i) PL pS1292-LRRK2, (ii) pT73-Rab10 western blot and (iii) PL pT73(Rab10)-Rab10, each of which yielded similar results.

In our studies, treating cells with a physiological concentration of H_2_O_2_ led to a rapid increase in LRRK2 activation within 5-10 minutes, which was prevented by quenching with NAC. This suggests that LRRK2 itself may be redox-sensitive. Similarly, various cellular perturbations that promote LRRK2 activation, such as mitochondrial stress (rotenone), lysosomal dyshomeostasis (chloroquine), and vesicular trafficking deficits (monensin) have also been shown to result in redox damage (*21, 22, 35, 36, 52–54*). Here we demonstrated that these perturbations can stimulate 4-HNE accumulation and subsequent adduct formation with LRRK2 (***fig. S15)***.

The dynamics of the kinase domain have a strong influence on the kinase activation state of LRRK2 (*12, 13*). We found that Cys2024/2025 in the kinase activation loop are important residues for stimulated LRRK2 activation. Recent work by others has also suggested that these cysteine residues may be important for kinase activation (*37*); however, they used *in vitro* kinase assays and experiments (*37*), whereas we maintained an endogenous cellular context and used multiple complementary assays to support our conclusions. We found that while the cysteine-to-alanine mutations preserved basal kinase activity, stimulated LRRK2 activity was completely prevented. The fact that LRRK2 bearing these cysteine-to-alanine mutations had little, if any, propensity to form adducts with 4-HNE suggests that 4-HNE preferentially targets these solvent-exposed Cys2024/2025 residues. It is important to note that the Cys2024/2025 to Ala mutants are not “kinase-dead”; they retain normal basal activity but simply cannot be further activated. A previous study postulated that the development of LRRK2 kinase inhibitors directed towards Cys2024 and Cys2025 may hold therapeutic benefit (*55*). Our study supports this hypothesis and indicates these residues are critical for LRRK2 hyperactivation. It is unlikely that redox stress causes Cys2024/2025 to form an intramolecular disulfide bridge in monomeric LRRK2, as disulfide bridge formation between vicinal cysteine residues is an extremely rare biological phenomenon (*56*). Whether intermolecular disulfide bonds involving these cysteine residues might occur in higher-order oligomeric species of LRRK2 (*57*) is unknown.

LRRK2 activation has been shown to be associated with damaged membranes, especially in the endolysosomal system (*20–23, 35*). As noted, endolysosomal impairment and mitochondrial stress lead to the formation of the lipid peroxidation product 4-HNE. This raises the possibility that this reactive aldehyde may be important in LRRK2 activation and might provide a link between membrane damage and LRRK2 activation. Given that 4-HNE is known to activate other kinases (*25–28*), there is precedent for this idea. Indeed, we found that treatment of cells with exogenous 4-HNE led to dose-dependent increases in LRRK2-4-HNE adduct formation and LRRK2 kinase activation – with a strong linear correlation between the two. Similarly, other LRRK2 activating stimuli (rotenone, chloroquine, monensin, and H_2_O_2_), which lead to the endogenous accumulation of 4-HNE, also resulted in LRRK2-4-HNE adduct formation. Interestingly, Rab GTPases, which have been shown to activate LRRK2 (*58, 59*), may also be modified by 4-HNE (*60*). Furthermore, STING pathway activation, which leads to lipid hydroperoxidation (*61*), plays a role in the activation of LRRK2 at damaged lysosomes (*23*). As such, 4-HNE formation appears to be a convergent event shared by these seemingly disparate LRRK2 activating stimuli. Thus, 4-HNE may be a final common regulator of LRRK2 kinase activation.

Levels of 4-HNE and 4-HNE protein adducts are elevated in PD patients compared to healthy controls (*46, 62*). 4-HNE can be formed nonenzymatically or enzymatically through the 15-LO pathway (*29, 30*). Therefore, we explored whether 15-LO activity regulates 4-HNE production, LRRK2–4-HNE adduct formation, and LRRK2 kinase activation. Remarkably, 15-LO knockout or 15-LO inhibition with CU-12991 potently blocked rotenone-stimulated 4-HNE production, LRRK2-4-HNE adducts, and LRRK2 kinase hyperactivation but did not inhibit basal kinase activity. Moreover, the 15-LO specific metabolite 15(S)-HpETE, which undergoes subsequent intramolecular rearrangement to form 4-HNE, dose-dependently increased LRRK2 activity. Importantly, we demonstrated in patient-derived iPD and G2019S PD LCLs that there are elevated basal 4-HNE levels and LRRK2 kinase activity that could be reduced to healthy control levels by inhibiting 15-LO with CU-12991. Thus, we conclude that the specific end-product of 15-LO activity, 4-HNE, is sufficient for endogenous pathologic LRRK2 activity (***fig. S15)***.

Normally, 15-LO prefers to hydroperoxidate free fatty acids, not those incorporated into membranes. However, its substrate specificity is regulated by interaction with PEBP1 (*48*). When 15-LO is in complex with PEBP1, it acquires the ability to oxygenate membrane phospholipids (*48*), which ultimately results in the formation of 4-HNE. In this regard, we found that rotenone led to an increase in the association of 15-LO with PEBP1 *in vitro* and *in vivo*. We found that stimuli that increase endogenous WT LRRK2 kinase activity also (i) enhance the PEBP1–15-LO interaction, (ii) increase 4-HNE production, and (iii) cause LRRK2-4-HNE adducts to form (***fig. S15)***. Strikingly, this association was observed in both G2019S PD and iPD patient-derived samples, confirming the relevance of this mechanism to human disease. These data suggest that the increase in 15-LO–PEBP1 interaction leads to elevated 4-HNE levels at specific membranes, which in turn promotes the hyperactivation of LRRK2. The fact that this sequence of events is demonstratable in G2019S PD and iPD patient-derived samples provides strong evidence for the 15-LO pathway as a key contributor to PD pathogenesis through the regulation of LRRK2 activity.

Our study has some limitations. We acknowledge that while our data strongly implicates cysteine residues 2024 and 2025 in the redox-dependent activation of LRRK2, we did not show directly that they undergo a specific 4-HNE post-translational modification. Additionally, we have not determined the subcellular localization of LRRK2 activation in response to these stimuli. It will be important to determine if LRRK2 activation and LRRK2–4-HNE interaction occur in one specific cellular compartment, or if it is dependent on the location of the inciting insult (*e.g.*, lysosomes, mitochondria).

In summary, we have defined a new mechanism of cellular perturbation-induced LRRK2 kinase activation. Our data demonstrate that LRRK2 is a redox-sensitive kinase and Cys2024/2025 are critical for stress-induced stimulation of kinase activity. Moreover, we found that seemingly disparate LRRK2 activating stimuli (chloroquine, H_2_O_2_, monensin, and rotenone), each with a different proximal mechanism of action, activate LRRK2 through a convergent redox-dependent mechanism that is mediated by the lipid hydroperoxidation end-product, 4-HNE, which forms adducts with LRRK2. Further, 15-LO, the enzymatic source of 4-HNE, was identified as the upstream mediator of stimulated LRRK2 kinase activation (***fig. S15)***. These results suggest that 15-LO inhibitors or drugs that otherwise reduce lipid hydroperoxidation may provide a new therapeutic avenue to target the LRRK2 pathway in PD. Moreover, the fact that inhibition of 15-LO prevented stimulus-induced LRRK2 kinase hyperactivation in cell lines, and disease-associated hyperactivity in LCLs, without impacting basal kinase activity, may allow therapeutic modulation of LRRK2 while avoiding on-target liabilities.

## MATERIALS AND METHODS

### Study Design

This study was designed to explore the mechanism(s) that drive endogenous WT LRRK2 kinase activation. To this end we used CRISPR/Cas9 gene-edited HEK293 cells, CRISPR/Cas9 gene-edited HAP1 cells, as well as RAW264.7 macrophages, patient-derived LCLs, and an *in vivo* rotenone rat model of PD to explore what physiological players regulate LRRK2 activation. To enhance scientific rigor, we generally used orthogonal approaches to support each conclusion. Furthermore, as target validation of 15-LO, specifically, 15-LO derived 4-HNE, being upstream LRRK2 activation, we employed a specific and highly potent pharmacological inhibitor and genetic knockout approaches to support key findings. To ensure clinical relevance and translational potential, key results were confirmed in patient-derived LCLs and in an *in vivo* model of PD. All *in vitro* experiments were replicated independently at least three times and performed by multiple experimenters. *In vivo* experiments were conducted using a single cohort of rats that were assigned randomly to treatment group.

### Cell Culture

HEK293 cells and RAW264.7 macrophages were cultured in DMEM/F12 (cat:11320082, Thermo Fisher) supplemented with 10% fetal bovine serum (FBS) and penicillin-streptomycin (pen-strep). 15-LO^WT/WT^ and CRISPR/Cas9 gene-edited 15-LO^-/-^ HAP1 cells were purchased from Horizon (HZGHC007334c00) and cultured in IMDM (cat: 30-2005, ATCC) supplemented with 10% FBS and pen-strep. Cells were maintained at 37°C and 5% CO_2._ CRISPR/Cas9 gene-edited HEK293 cells were generated as previously described (*9*). LRRK2^-/-^ (sc-6004) and LRRK2^WT/WT^ (parental) (sc-6003) RAW264.7 macrophages were obtained from ATCC. Cell passage number did not exceed 20 for HEK293 or RAW264.7 macrophages. Cells were plated on poly-d-lysine coated coverslips in a 24-well plate or 16-well Nunc Lab Tek chamber slides (cat: 178599, Thermo Fisher). Details for reagents used for cell treatments can be found in Table S1. For H_2_O_2_, 4-HNE, 12(S)-HpETE, and 15(S)-HpETE treatments, DMEM/F12 supplemented with 5% fetal bovine serum (FBS) and penicillin-streptomycin (pen-strep) was used.

Patient-derived lymphoblastoid cell lines were obtained from the NINDS Coriell Biorepository through an MTA and sample ID numbers may be found in Table S2. Cells were maintained in RPMI medium GlutaMAX supplement (cat: 61870127, Thermo Fisher Scientific) with 15% FBS and pen-strep. Cells were plated on poly-d-lysine coated coverslips in a 24-well plate or 16-well Nunc Lab Tek chamber slides (cat: 178599, Thermo Fisher) and attached by light centrifugation at 690 x g at room temperature. Cells were passaged every 3 days and did not exceed 20 passages.

### CRISPR/Cas9 gene edited HEK293 cells (C2024A, C2025A, C2024A/C2025A)

LRRK2^C2024A/C2024A^, LRRK2^C2025A/C2025A^ and double cysteine mutants were created in-house. A guide RNA targeting exon 41 of the LRRK2 gene (5’-CTCAGTACTGCTGTAGAATG-3’) was commercially synthesized (ThermoFisher) and used to form ribonucleoprotein (RNP) complexes when combined with recombinant Cas9 protein (New England Biolabs). Ribonucleoprotein complexes were formed using the method of Kouranova^49^. RNP complexes were formed by incubating the recombinant Cas9 protein and guide RNA at 1:1 mass ratio (1: 4.6 molar ratio of the Cas9 protein to sgRNA) at 37°C for 5 min. RNPs were delivered to HEK293 cells using Transit-X2 (Mirius) transfection reagent as described by the manufacturer. Single stranded repair templates specific to the C2024 or C2025 residues were included in the transfection reactions to facilitate the cysteine to alanine conversion by homologous recombination. The double mutant C2024A/C2025A was generated in a similar fashion using the repair template specific to the double mutation. Transfected cells were collected and enriched by FACS sorting. Sorted cells were grown and expanded for PCR and DNA sequencing analyses. All clones generated by these methods were sequenced on both strands to confirm their identity. Protein expression – or lack thereof – was confirmed by western blotting. Using the same technology, LRRK2^G2019S/G2019S^, LRRK2^R1441G/R1441G^, LRRK2^-/-^ cells were created. Details for CRISPR/Cas9 reagents can be found in Table S3.

### Dihydroethidium (DHE) Staining

DHE staining was performed as described previously (*44*). A 3 mM stock solution of DHE (cat: D11347, Thermo Fisher Scientific) was reconstituted in DMSO and stored at -20°C in single use aliquots. Prior to use, the DHE stock solution was thawed, protected from light, and diluted in cell culture media to reach a final concentration of 3μM DHE. Old media was aspirated and replaced with the fresh media containing 3μM DHE. Cells were incubated for 20 minutes at 37°C and 5% CO_2_, protected from light. Cells were then gently washed with phosphate buffer saline (PBS) at room temperature for 5 minutes and then fixed with 4% paraformaldehyde (PFA) (cat:15710-S, Electron Microscopy Science) for 20 minutes. Following fixation, cells underwent three 10-minute washes in PBS. Coverslips were mounted on glass slides (cat: 22-037-246, Fisher Scientific) using gelvatol mounting medium (see (*34*) for recipe).

### Immunofluorescence

Cells were fixed in 4% paraformaldehyde in phosphate buffer saline (PBS) for 20 minutes at room temperature, followed by three 10-minute washes in PBS on a shaker. Cells were blocked and permeabilized in 10% normal donkey serum (NDS) + 0.02% triton-x in PBS for 1 hour at room temperature on a shaker. After blocking/permeabilization, cells underwent three washes in PBS and then were incubated in primary antibody (Table S4, for antibody details) in PBS containing 1% NDS overnight at 4°C on a shaker. The following day, cells underwent three 10-minute washes in PBS and were incubated in secondary antibody in PBS containing 1% NDS for one hour at room temperature on a shaker protected from light. Cells underwent three 10-minute washes in PBS and stained with DAPI for two minutes, followed by two quick washes with PBS. Coverslips were mounted on glass slides (cat: 22-037-246, Fisher Scientific) using gelvatol mounting medium.

### Proximity Ligation Assay

#### Cells (*Figures 1 and 3*)

Proximity ligation assay (PLA) was performed on an individual coverslip in 24 well plates using Duolink In Situ PLA reagents (DUO92002, DUO920024, DUO92007 (orange) or DUO92014 (green), Sigma Aldrich). Cells were fixed, blocked/permeabilized, and incubated in primary antibody solution as above and as previously described (*34*). Following overnight primary antibody incubation (Table S4, for antibody details), cells underwent three 10-minute washes in PBS. After washes, PLA was performed as described in (*34*). PLA incubation steps took place in a humidified incubator at 37°C with gentle agitation taking care that samples did not dry out. For details of other products used see (*34*).

#### Cells

Proximity ligation assay (PLA) was performed in 16-well Nunc Lab Tek chamber slides (cat: 178599, Thermo Fisher) using a NaveniFlex MR Kit (cat: NF.MR.100, Navinci). Cells were fixed, blocked/permeabilized, and incubated in primary antibody solution as above and as previously described (*34*). Following overnight primary antibody incubation (Table S4, for antibody details), cells underwent three 10-minute washes in PBS. After washes, PLA was performed according to manufacturer’s protocol. PLA incubation steps took place in a humidified incubator at 37°C with gentle agitation taking care that samples did not dry out. For details of other products used see (*34*).

#### Rat brain tissue

PLA was performed in blocked and permeabilized brain tissue as previously described (*34*) using NaveniFlex Tissue MR Red (NT.MR.100.RED). Free floating tissue sections were blocked and permeabilized in 1% triton-x (in PBS) containing 10% NDS for one hour at room temperature on a shaker. Sections were then washed three times (10 minutes per wash) and incubated in primary antibody solution (1% NDS in PBS) to label tyrosine hydroxylase (Table S4, for antibody details) overnight at 4°C on a shaker. The following day the tissue sections underwent three 10-minute washes in PBS and were incubated in secondary antibody in PBS containing 1% NDS for one hour at room temperature on a shaker protected from light. After secondary antibody labeling, tissue sections were incubated overnight 4°C on a shaker in primary antibody solution containing the antibodies for PLA (Table S4, for antibody details). Next day, tissue sections underwent three 10-minute washes in PBS and then were mounted on glass slides (cat: 22-037-246, Fisher Scientific). See (*34*) for detailed description and video for mounting of tissue prior to PLA. PLA was performed following the manufacturer’s protocol. PLA incubation steps took place at 37°C ensuring that samples did not dry out. After PLA was completed, sections were covered with a glass coverslip with gelvatol mounting medium.

### Animals

All experiments utilizing animals were approved by the Institutional Animal Care and Use Committee of the University of Pittsburgh. Male Lewis rats (8-9 months old) received a single daily interperitoneal (i.p.) injection of 2.8 mg/kg of rotenone resuspended in 2% DMSO, 98% miglyol 812 N as previously described (*45, 63*) until they reached behavioral endpoint (7-10 injections). Animals were euthanized using 0.3 mg/kg pentobarbital, followed by transcardial perfusion. Brains were removed and postfixed in 4% paraformaldehyde for 24-h before placing them 30% sucrose. Free-floating sections (35μm) were collected using a microtome and stored in cyroprotectant at −20 °C until use.

### Cell lysis and western blot

Cells were collected on ice in 1X Cell Signaling lysis buffer (cat:9803, Cell Signaling) containing 1:33 Halt Phosphatase Inhibitors (cat:78427, Thermo Fisher Scientific) and 1:100 Halt Protease and Phosphatase Inhibitors (cat:78446, Thermo Fisher Scientific). Cells were lysed using a freeze-thaw lysis method where cells were incubated in pulverized dry ice with ethanol for three minutes followed by a warm water bath for three minutes and was repeated three times. Lysates were clarified by centrifugation at 12,000 rpm for 10 minutes 4°C. Supernatant of the cell lysates were collected, and protein quantified by Pierce BCA kit. 25-45μg of samples (dependent on target) was mixed with 4X NuPAGE LDS Sample Buffer (supplemented with 5% β-mercaptoethanol and 10% glycerol). Samples were run in duplicates on 4-12% NuPAGE or 10% NuPAGE Bis-Tris midi gels with MOPS Running Buffer at 120 Volts for eight minutes followed by 200 Volts for 45 minutes. At the end of electrophoresis, samples were transferred onto nitrocellulose membranes (0.2μm) at 100 Volts for 60 minutes on ice in prechilled transfer buffer (25mM Tris-Base, 198mM Glycine, 20% methanol v/v). Transferred membranes were blocked in Fish Serum Blocking Buffer (cat:37527, Thermo Scientific) diluted in PBS (1:1 ratio) for one hour at room temperature on a shaker. Membranes were incubated in primary antibodies (Table S4, for antibody details) diluted in Fish Serum Blocking Buffer/PBS overnight at 4°C on a shaker. The following day, membranes were washed in PBS three times 10 minutes each and incubated in LiCor secondary antibodies (Table S4, for antibody details). diluted in Fish Serum Blocking Buffer/PBS for 2 hours at room temperature on a shaker. Membranes were washed three times in PBS for a duration of 10 minutes each and imaged on an Odyssey LiCor CLx imaging system. Analysis was performed using Image Studio software.

### Confocal microscopy and analysis

Images were acquired on a Nikon Eclipse Ti2 Resonance Scan Spectral Confocal Microscope at 60X magnification using Nyquist criterion and resonant scanning. Laser parameters were set up on positive and negative controls (e.g. primary antibody delete) to ensure there was no pixel saturation. For quantitative comparisons all imaging parameters (e.g. laser power, gain, pinhole) were held constant across an experiment. Confocal images were analyzed using Nikon NIS-Elements Advanced Research software. For DHE fluorescence intensity analysis, regions of interest were drawn around the signal on a per cell basis to obtain the fluorescence intensity. For all other analyses, a binary layer was created using threshold parameters for each experiment and used to identify each puncta or ‘object’.

For Figures 1 and 3 images were acquired on Olympus BX61 confocal microscope at 60X magnification (UPlansApo 60X) and Fluoview 1000 software (Melville, NY). Imaging parameters were monitored to ensure that images were above background level and below pixel saturation. For quantitative comparisons between groups, all imaging parameters (e.g. laser power, exposure, and pinhole) were held constant across. For PL pS1292-LRRK2 (Figures 1 and 3, Sigma) intensity analysis, regions of interest were drawn around the signal on a per cell basis to obtain the fluorescence intensity. LRRK2^-/-^ cells were used as assay controls.

#### Colocalization Analysis

Our protocol is similar to that of the University of Pittsburgh Center for Biologic Imaging (*64*). Images are acquired on a Nikon A1R HD25 High-Definition Resonant Scanning Confocal System with a 60X objective (1.4 NA) using Nyquist criterion. Exposure time of each channel is kept consistent throughout sampling. Images were analyzed using NIS Elements 5.2 software. Separate binary layers are created for the 15-LO protein immunoreactivity and PEBP1 signal. The intersection tool is then used to identify the third binary layer, the “colocalization” mask, which corresponds to the overlapping immunoreactivity signals. The threshold for each channel is kept consistent throughout the analysis per experiment. In this way, the percent of 15-LO immunoreactivity (based on the binary mask) that colocalizes with PEBP1 signal is derived.

### Statistical Analysis

Results are presented as mean ± SEM and are derived from 3 - 5 independent experiments. For simple comparisons of two experimental conditions, two-tailed unpaired *t-*tests were used. Where variances were unequal Welch’s correction was used. For comparisons of multiple experimental conditions, one-way or two-way ANOVA was used, and if significant overall, post hoc corrections (with Tukey tests) for multiple comparisons were made. P-values less than 0.05 were considered significant.

## Acknowledgments

Figure S15 was made using Biorender (Agreement #: FU26XHYZDS)

## Funding

National Institutes of Health grant R56 NS131137-01 (JTG)

The friends and family of Sean Logan (JTG)

The Blechman Foundation for Parkinson’s Research (JTG)

The American Parkinson Disease Association Center for Advanced Research at the University of Pittsburgh (JTG)

The Michael J Fox Foundation Pritzker Prize (JTG)

Commonwealth of Pennsylvania Grant #601457

## Author contributions

Conceptualization: MTK, TGH, SAP, WDS, JTG Methodology: MTK, SAP, WDS, TGH, JTG

Investigation: MTK, EKH, JW, WGW, SC, KF, MF, RDM, AK

Visualization: MTK, SAP, WDS, JTG

Funding acquisition: JTG

Project administration: MTK, JTG

Supervision: JTG

Writing – original draft: MTK, JTG

Writing – review & editing: MTK, EMR, TGH, SAP, WDS, JTG

## Competing interests

WDS and SAP are full-time employees of Acurex Biosciences and also own stock in the company. AK is a paid consultant of Acurex Biosciences.

## Data and materials availability

All data are available in the main text or the supplementary materials.

## Notes

### Summary of Updates

This version corrected the author order and supplemental materials were uploaded.

