## Supplemental for "15-Lipoxygenase-Mediated Lipid Peroxidation Regulates LRRK2 Kinase Activity"

**Affiliations:**
[1]Pittsburgh Institute for Neurodegenerative Diseases, University of Pittsburgh School of Medicine, Pittsburgh, Pennsylvania.
[2]Department of Neurology, University of Pittsburgh School of Medicine, Pittsburgh, Pennsylvania.
[3]Department of Pharmacology and Chemical Biology, University of Pittsburgh School of Medicine, Pittsburgh, Pennsylvania.
[4]Acurex Biosciences, San Carlos, California.


**List of Supplementary Materials**


Figure S1: N-acetyl cysteine prevents rotenone induced Rab10 phosphorylation
Figure S2: PL pT73(Rab10)-Rab10 validation
Figure S3: Chloroquine and monensin cause ROS production and lipid peroxidation
Figure S4: C2024A or C2025A mutations do not affect baseline kinase activity
Figure S5: C2024A or C2025A prevent rotenone induced ROS production
Figure S6: Exogenous 4-HNE forms adducts with LRRK2
Figure S7: Exogenous 4-HNE induces kinase activation
Figure S8: Correlation between 4-HNE induced kinase activity and LRRK2-4HNE interaction
Figure S9: Rotenone induces endogenous LRRK2-4HNE adduct formation
Figure S10: LRRK2 activating stimuli promote the endogenous LRRK2-4HNE adduct formation
Figure S11: Structure and $IC_{50}$ of CU-12991
Figure S12: 15-LO$^{-/-}$ prevents rotenone induced 4-HNE accumulation and Rab10 phosphorylation
Figure S13: CU-12991 does not interfere with 4-HNE
Figure S14: CU-12991 prevents stimulated Rab10 phosphorylation
Figure S15: 4-HNE mediated LRRK2 activation

Table S1 Reagents used for cell culture treatments
Table S2 Lymphoblastoid cell line information
Table S3 CRISPR/Cas9 reagents
Table S4 List of antibodies used for immunofluorescence, proximity ligation assay, and western blot

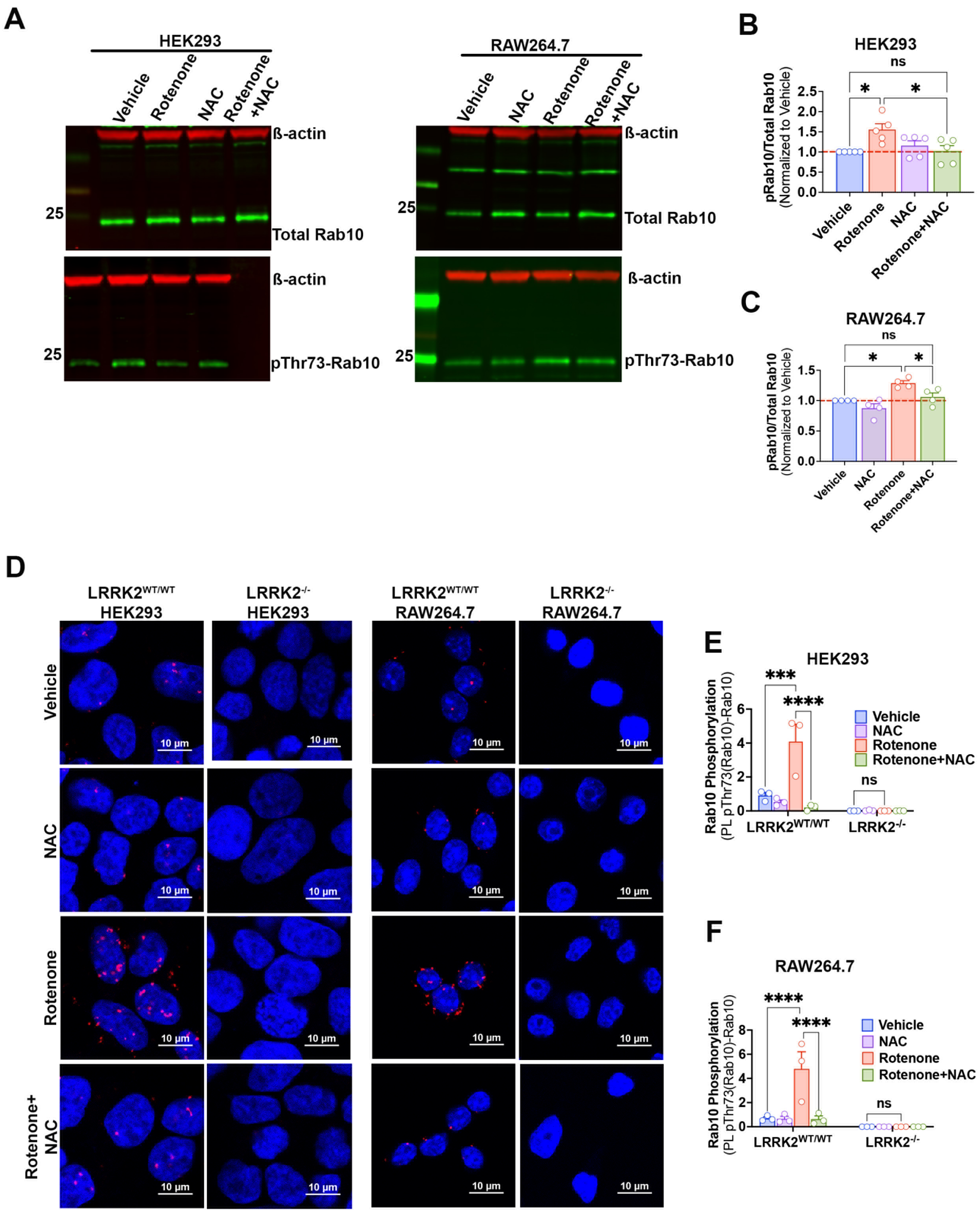


**Fig S1. N-acetylcysteine prevents rotenone induced Rab10 phosphorylation**. (**A**) Representative western blots for total Rab10 (top) and pThr73-Rab10 (bottom) in LRRK2$^{WT/WT}$ HEK293 cells (left) and RAW264.7 macrophages (right). Rotenone elicited an increase in pT73-Rab10/Total Rab10 compared to vehicle in both cell types. Cotreatment with NAC prevented the rotenone induced increase pT73-Rab10/Total Rab10. Total Rab10 levels were not changed. (**B**) Quantification of pT73-Rab10/Total Rab10 normalized to LRRK2$^{WT/WT}$ vehicle HEK293 cells. Each symbol represents the average of pRab10/Total Rab10 obtained from two technical replicates per treatment group for each of five independent experiments. Statistical testing by two-way ANOVA with Tukey correction. * denotes $p<0.05$; ns denotes not significant. (**C**) Quantification of pRab10/Total Rab10 normalized to LRRK2$^{WT/WT}$ vehicle

RAW264.7 macrophages. Each symbol represents the average of pT73-Rab10/Total Rab10 obtained from two technical replicates per treatment group for each of four independent experiments. Statistical testing by two-way ANOVA with Tukey correction. (**D**) Proximity ligation (PL) assay between pT73-Rab10 and total Rab10 (PL pT73(Rab10)–Rab10) was used to assess levels of Rab10 phosphorylation. In LRRK2$^{WT/WT}$ HEK293 and RAW264.7 macrophages, rotenone elicited an increase in PL pT73(Rab10)–Rab10 signal relative to vehicle. This was not observed in cells cotreated with NAC or in LRRK2$^{-/-}$ cells. (**E**) Quantification of Rab10 phosphorylation by PL pT73(Rab10)–Rab10 Objects/DAPI in HEK293 cells. Each symbol represents the average number of PL pT73(Rab10)–Rab10 Objects/DAPI obtained from 100-150 cells per treatment group for each of three independent experiments. Statistical testing by two-way ANOVA with post-hoc Tukey correction. (**F**) Quantification of Rab10 phosphorylation by PL pT73(Rab10)–Rab10 Objects/DAPI in RAW264.7 macrophages. Each symbol represents the average number of PL pT73(Rab10)–Rab10 Objects/DAPI obtained from 100-150 cells per treatment group for three independent experiments. Statistical testing by two-way ANOVA with post-hoc Tukey correction. * denotes $p<0.05$; *** denotes $p<0.0005$; **** denotes $p<0.0001$; ns denotes not significant.

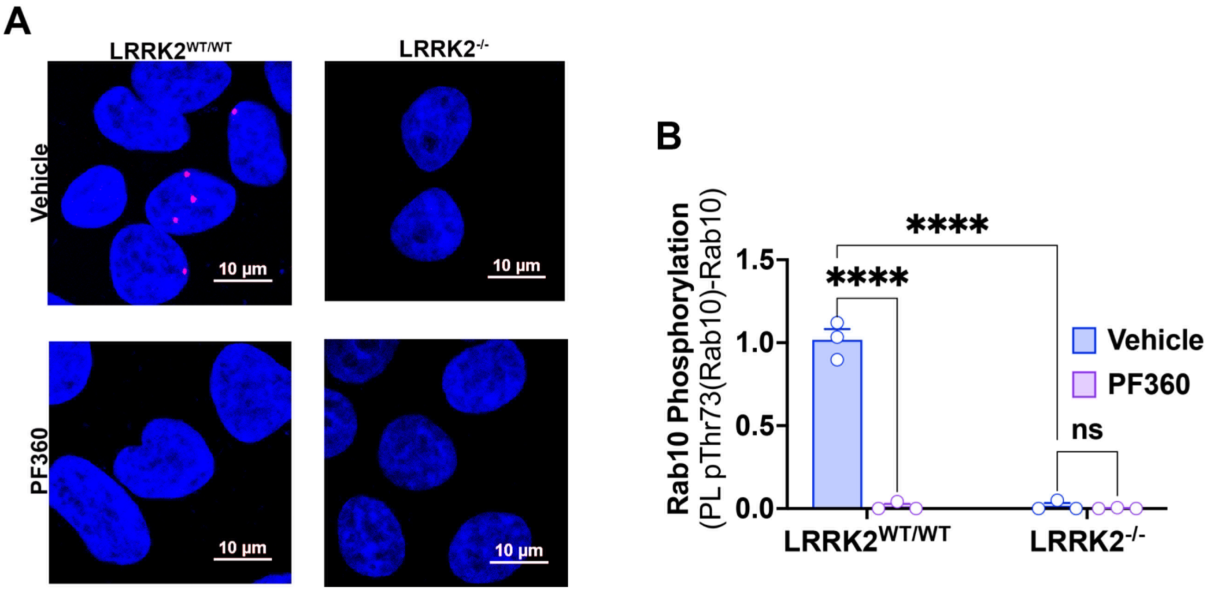


**Fig S2. PL pT73(Rab10)-Rab10 Validation*.*** (**A**) Validation of PL between pT73-Rab10 and total Rab10 (PL pT73(Rab10)–Rab10) to detect endogenous Rab10 phosphorylation in LRRK2$^{WT/WT}$ and LRRK2$^{-/-}$ HEK293 cells. At baseline, there was PL signal in LRRK2$^{WT/WT}$ which was significantly decreased in presence of the LRRK2 kinase inhibitor PF360. In contrast, there was no signal in LRRK2$^{-/-}$ HEK293 cells. (**B**) Quantification of Rab10 phosphorylation by PL pT73(Rab10)–Rab10 Objects/DAPI in HEK293 cells. Each symbol represents the average number of PL pT73(Rab10)–Rab10 Objects/DAPI obtained from 100 - 150 cells per treatment group for three independent experiments. Statistical testing by two-way ANOVA with post-hoc Tukey correction; **** denotes $p<0.0001$; ns denotes not significant.

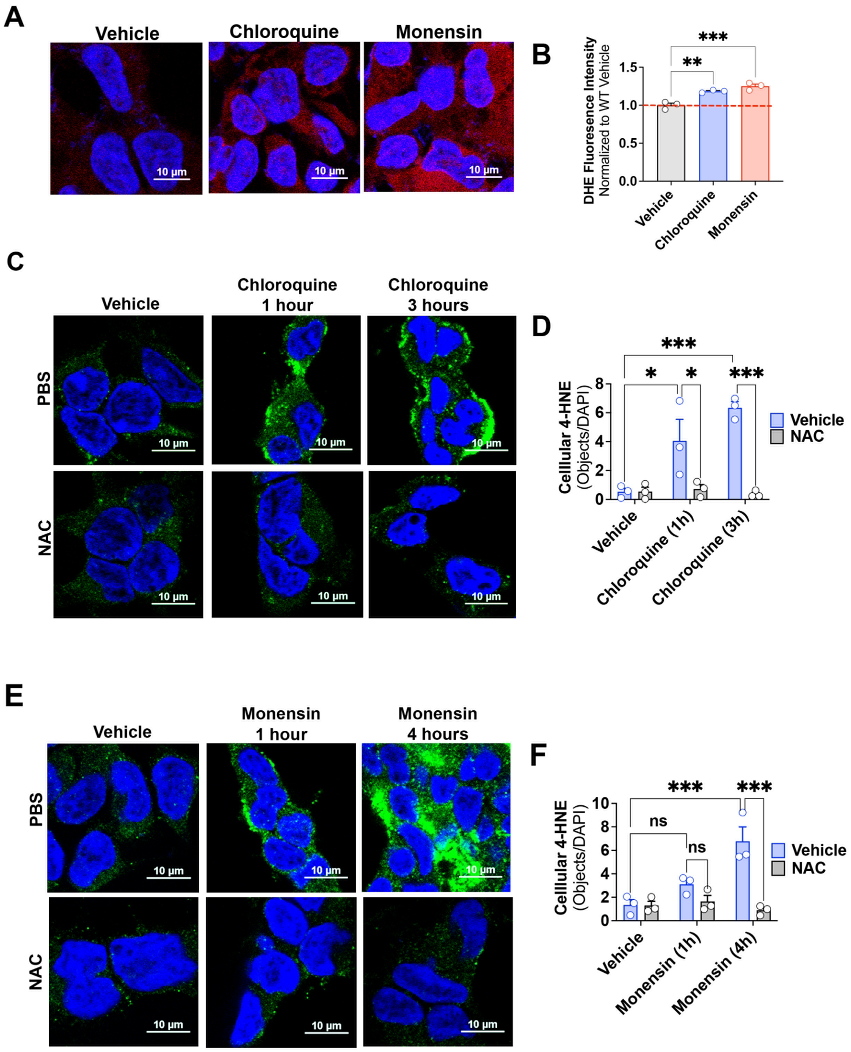


**Fig S3. Chloroquine and monensin cause ROS production and lipid peroxidation.** (**A**) ROS production was assessed by dihydroethidium (DHE) fluorescence (red). Chloroquine (100µM, 3 hours) and monensin (10µM, 4 hours) resulted in an increase in DHE signal compared to vehicle in LRRK2$^{WT/WT}$ HEK293 cells. (**B**) Quantification of normalized DHE signal. Each symbol represents the normalized average DHE signal measured from 100-150 cells per treatment group per experiment. N=3 independent experiments. Statistical testing by one-way ANOVA with post-hoc Tukey correction. (**C**) Immunofluorescence staining for 4-HNE (green) in LRRK2$^{WT/WT}$ HEK293 cells. LRRK2$^{WT/WT}$ HEK293 cells exposed to chloroquine had an increase in 4-HNE signal at 1 hour and 3 hours after treatment, which was prevented by co-treatment with NAC. (**D**) Quantification of cellular 4-HNE signal. Each symbol represents the average number of 4-HNE objects/DAPI values obtained from 100-150 cells per treatment group per experiment. N=3 independent experiments. Statistical testing by one-way ANOVA with post-hoc Tukey correction. (**E**) LRRK2$^{WT/WT}$ HEK293 cells treated with monensin displayed an increase in 4-HNE signal at 4 hours, which was prevented by NAC cotreatment. (**F**) Quantification of cellular 4-HNE signal. Each symbol represents the average number of 4-HNE objects/DAPI obtained from 100-150 cells per treatment group for each of three independent experiments. * denotes $p<0.05$; ** denotes $p<0.005$; *** denotes $p<0.001$; ns denotes not significant.

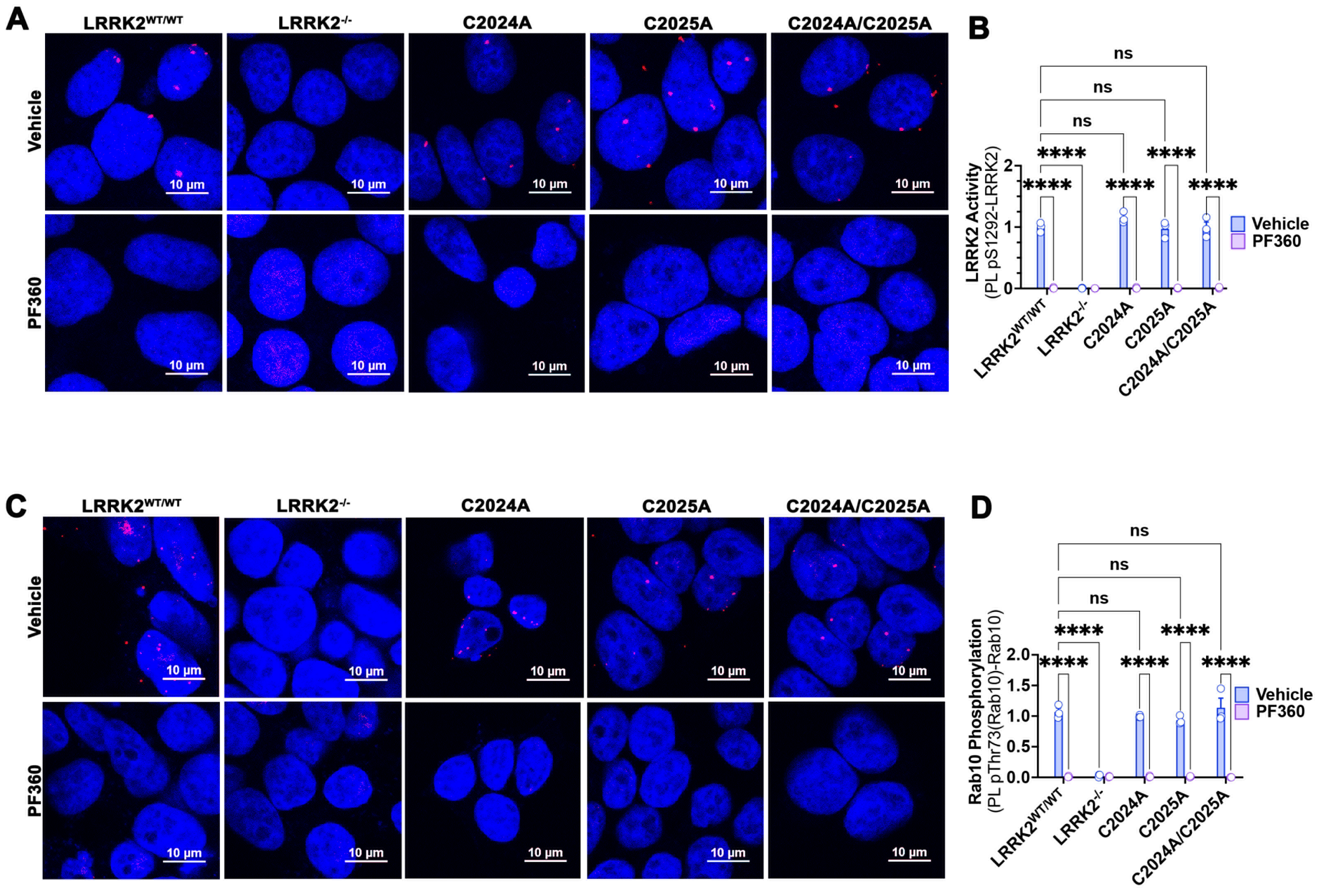


**Fig S4. C2024A or C2025A mutation do not affect baseline kinase activity. (A)** LRRK2 kinase activity was assessed by PL pS1292–LRRK2. Under vehicle conditions, C2024A, C2025A, and C2024A/C2025A had PL pS1292–LRRK2 signal (red dots) that was not different than LRRK2$^{WT/WT}$. Treatment with PF360 reduced PL pS1292-LRRK2 signal in LRRK2$^{WT/WT}$, C2024A, C2025A, and C2024A/C2025A. No signal was observed in LRRK2$^{-/-}$. (**B**) Quantification of LRRK2 activity by PL pS1292–LRRK2 Objects/DAPI. Each symbol represents the average number of PL pS1292-LRRK2 objects/DAPI obtained from 100-150 cells per treatment group for each of three independent experiments. Statistical testing by two-way ANOVA with post-hoc Tukey correction. (**C**) PL pT73(Rab10)–Rab10 was used to assess baseline Rab10 phosphorylation. Under vehicle conditions, C2024A, C2025A, and C2024A/C2025A PL pRab10–Rab10 signal (red dots) that was not different than LRRK2$^{WT/WT}$. Treatment with PF360 reduced PL pT73(Rab10)–Rab10 signal in LRRK2$^{WT/WT}$, C2024A, C2025A, and C2024A/C2025A. No signal was observed in LRRK2$^{-/-}$. (**B**) Quantification of Rab10 phosphorylation by PL pT73(Rab10)–Rab10 Objects/DAPI. Each symbol represents the average number PL pT73(Rab10)–Rab10 objects/DAPI obtained from 100-150 cells per treatment group for three independent experiments. Statistical testing by two-way ANOVA with post-hoc Tukey correction. **** denotes p<0.0001; ns denotes not significant.

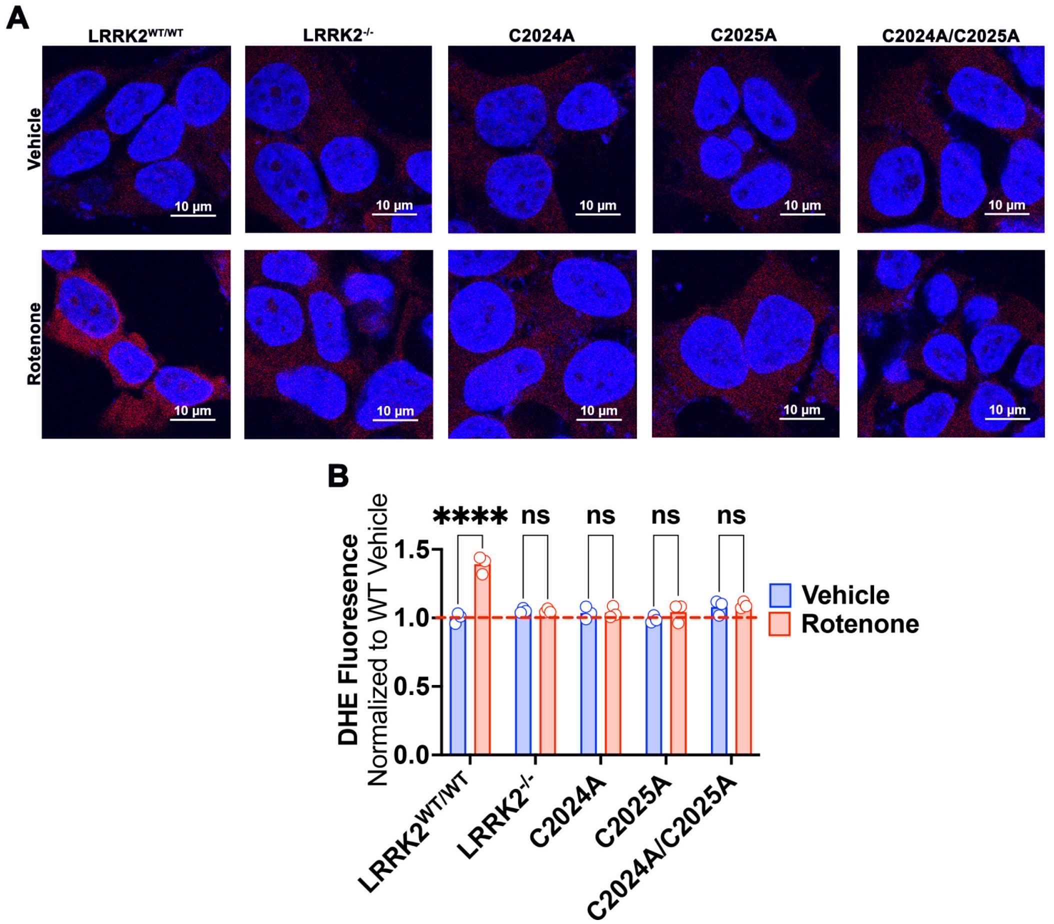


**Fig S5. C2024A or C2025A or double mutants prevent rotenone induced ROS production.** (**A**) ROS production was assessed by DHE fluorescence (red). Rotenone increased ROS in $LRRK2^{WT/WT}$ HEK293 cells but not in $LRRK2^{-/-}$, C2024A, C2025A, C2024A/C2025A cells. (**B**) Quantification of normalized DHE signal. Each symbol represents the normalized average DHE signal measured from 100-150 cells per treatment group for each of three independent experiments. Statistical testing by one-way ANOVA with post-hoc Tukey correction. **** denotes $p<0.0001$; ns denotes not significant.

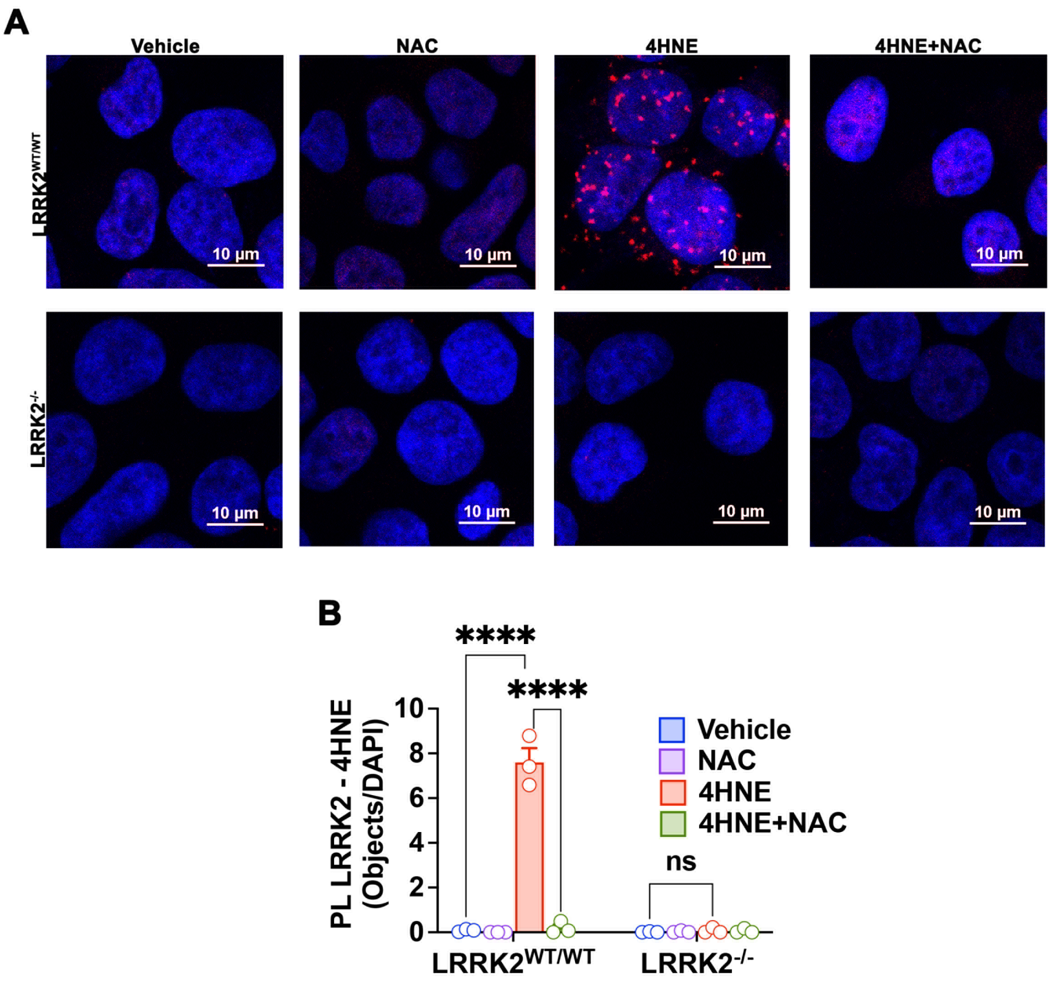


**Fig S6. Exogenous 4-HNE forms adducts with LRRK2.** (**A**) PL between LRRK2 and 4HNE (PL LRRK2–4HNE) was used as an index of LRRK2-4HNE adduct formation in LRRK2$^{WT/WT}$ and LRRK2$^{-/-}$ HEK293 cells. LRRK2$^{WT/WT}$ cells treated with 100 µM 4-HNE for 1 hour had strong PL LRRK2-4HNE signal compared to vehicle. This was prevented by NAC treatment and not observed in LRRK2$^{-/-}$ as an assay control. (**B**) Quantification of PL LRRK2-4HNE Objects/DAPI. Each symbol represents the average number of PL LRRK2–4HNE Objects/DAPI obtained from 100-150 cells per treatment group for each of three independent experiments. Statistical testing by two-way ANOVA with post-hoc Tukey correction. **** denotes $p<0.0001$; ns denotes not significant.

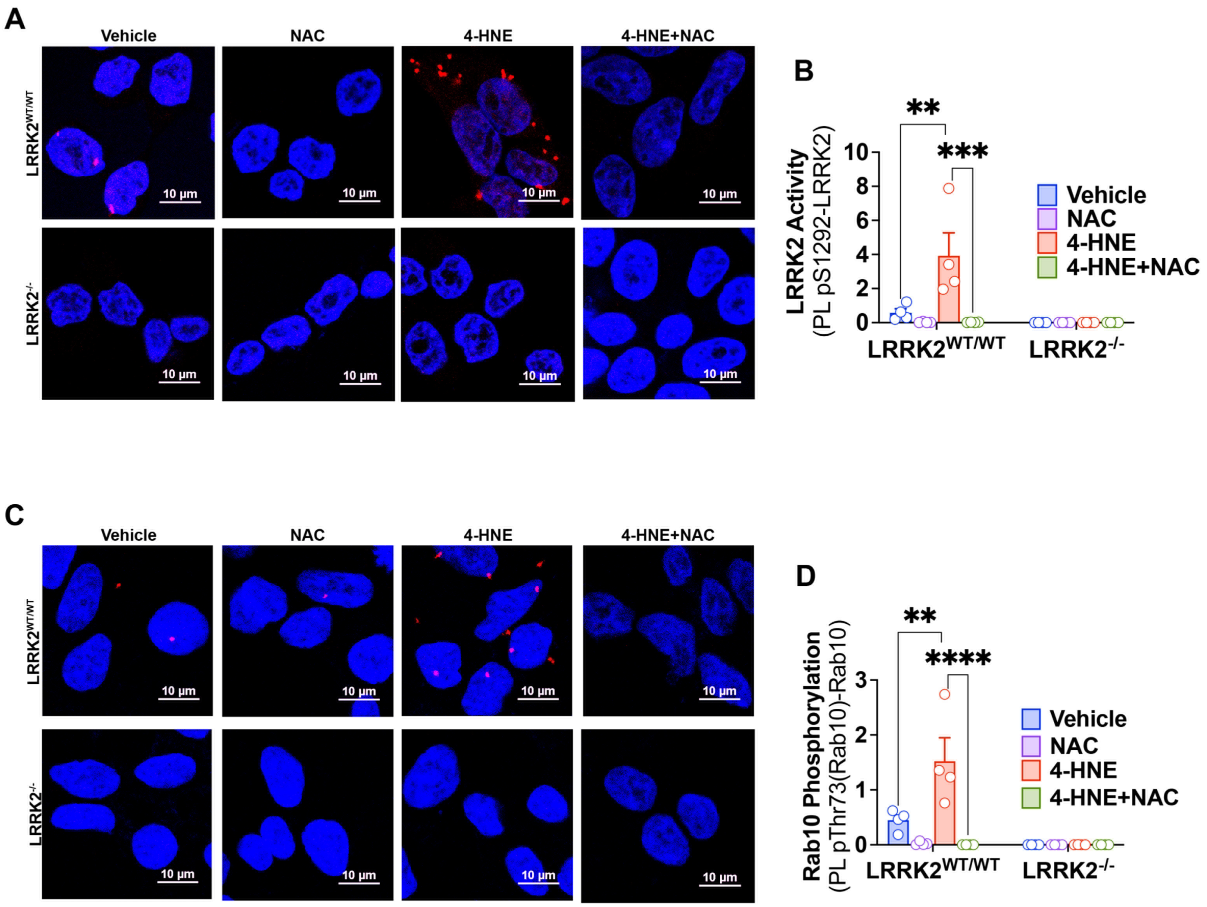


**Fig S7: Exogenous 4-HNE induces LRRK2 kinase activation.** (**A**) LRRK2 kinase activity was assessed by PL pS1292–LRRK2. LRRK2$^{WT/WT}$ HEK293 cells treated with 100μM 4-HNE had an increase in PL pS1292–LRRK2 signal (red dots) relative to vehicle, which was prevented by pretreatment with NAC. (**B**) Quantification of LRRK2 activity by PL pS1292–LRRK2 Objects/DAPI. Each symbol represents the average number of PL pS1292-LRRK2 objects/DAPI 100-150 cells per treatment group for each independent experiment; N=4 independent experiments. Statistical testing by two-way ANOVA with post-hoc Tukey correction. (**C**) PL pT73(Rab10)–Rab10 was used to assess 4-HNE induced Rab10 phosphorylation as a surrogate marker for LRRK2 kinase activity. LRRK2$^{WT/WT}$ HEK293 cells treated with 100μM 4HNE had an increase in PL pT73(Rab10)–Rab10 signal (red dots) relative to vehicle, which was prevented by pretreatment with NAC. (**B**) Quantification of Rab10 phosphorylation by PL pT73(Rab10)–Rab10 Objects/DAPI. Each symbol represents the average number PL pT73(Rab10)–Rab10 Objects/DAPI obtained from 100-150 cells per treatment group for each of four independent experiments. Statistical testing by two-way ANOVA with post-hoc Tukey correction. ** denotes p<0.005; *** denotes p<0.0005; **** denotes p<0.0001.

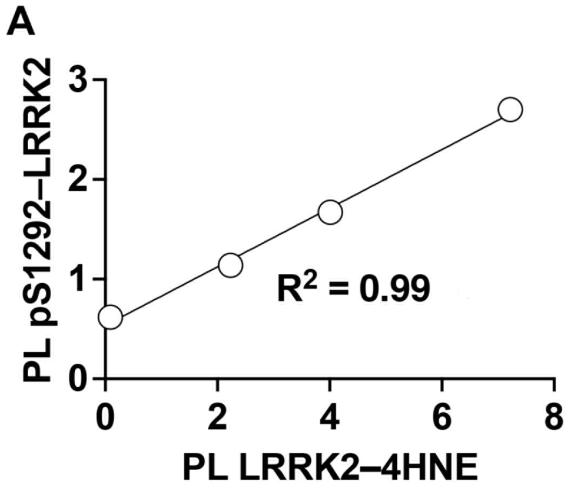


**Fig S8. Correlation between LRRK2-4HNE adduct formation and LRRK2 kinase activity.** LRRK2$^{WT/WT}$ HEK293 cells were treated with 4-HNE (10μM, 30μM, 100μM) and, in separate experiments, PL LRRK2-4HNE and PL pS1292-LRRK2 were measured as seen in Figures 4C and 4D. For each concentration of 4-HNE, mean values for PL LRRK2-4HNE and PL pS1292-LRRK2 were plotted and correlation analysis was performed. $R^2 = 0.996$; $p<0.005$.

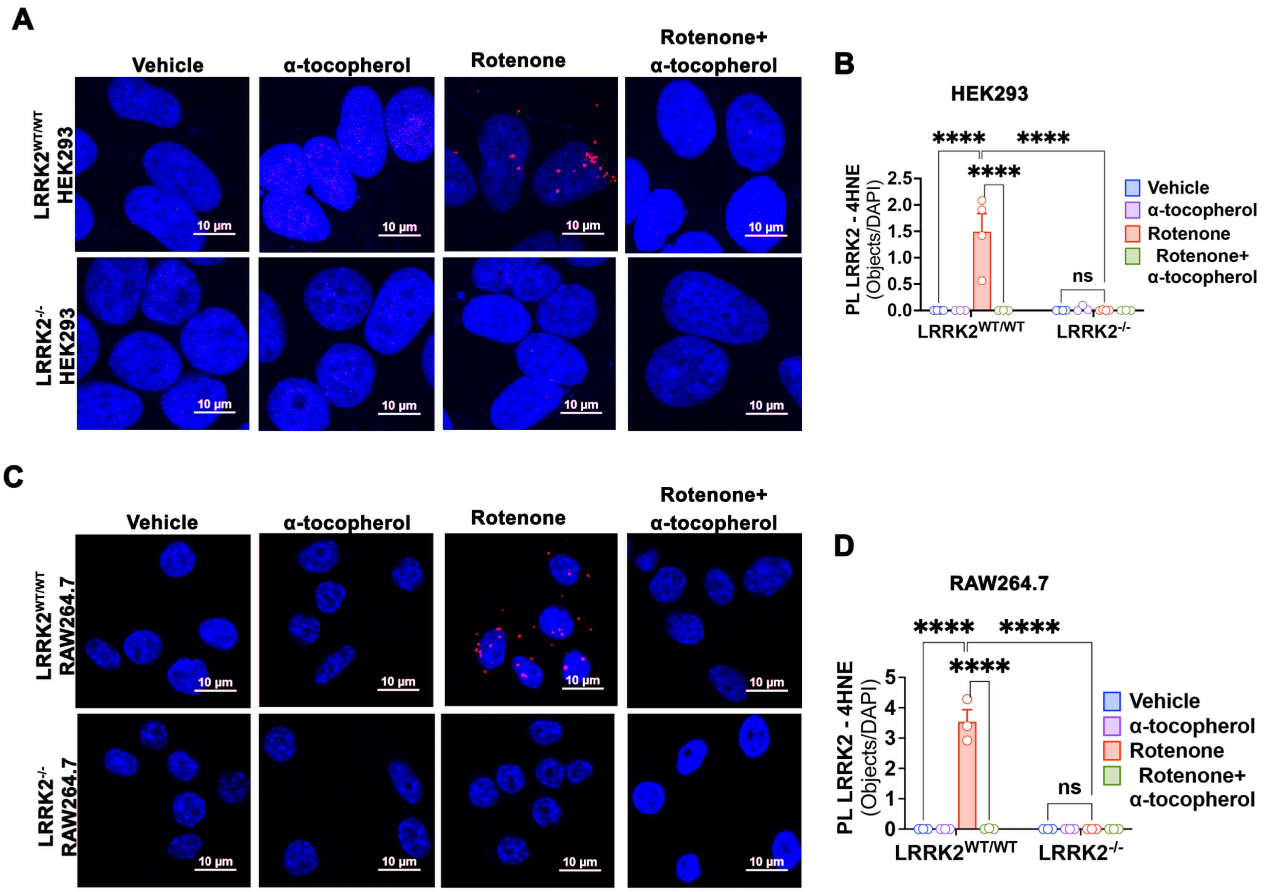


**Fig S9. Rotenone induces endogenous LRRK2-4HNE adduct formation.** Endogenous LRRK2 – 4-HNE adduct formation was assessed by PL LRRK2–4HNE in LRRK2$^{WT/WT}$ and LRRK2$^{-/-}$ (**A**) HEK293 and (**C**) RAW264.7 macrophages. Rotenone treatment increased PL LRRK2–4HNE signal (red dots) in LRRK2$^{WT/WT}$ (**A**) HEK293 and (**C**) RAW264.7 macrophages. Rotenone-induced PL LRRK2–4HNE signal was prevented by cotreatment with the lipid soluble antioxidant, $\alpha$-tocopherol. (**B**) Quantification of PL LRRK2–4HNE Objects/DAPI in HEK293 cells. (**D**) Quantification of PL LRRK2–4HNE Objects/DAPI in RAW264.7 macrophages. For each graph, each symbol represents the average number of PL LRRK2–4HNE Objects/DAPI obtained from 100-150 cells per treatment group for each of three independent experiments. Statistical testing by two-way ANOVA with post-hoc Tukey correction. **** denotes p<0.0001; ns denotes not significant.

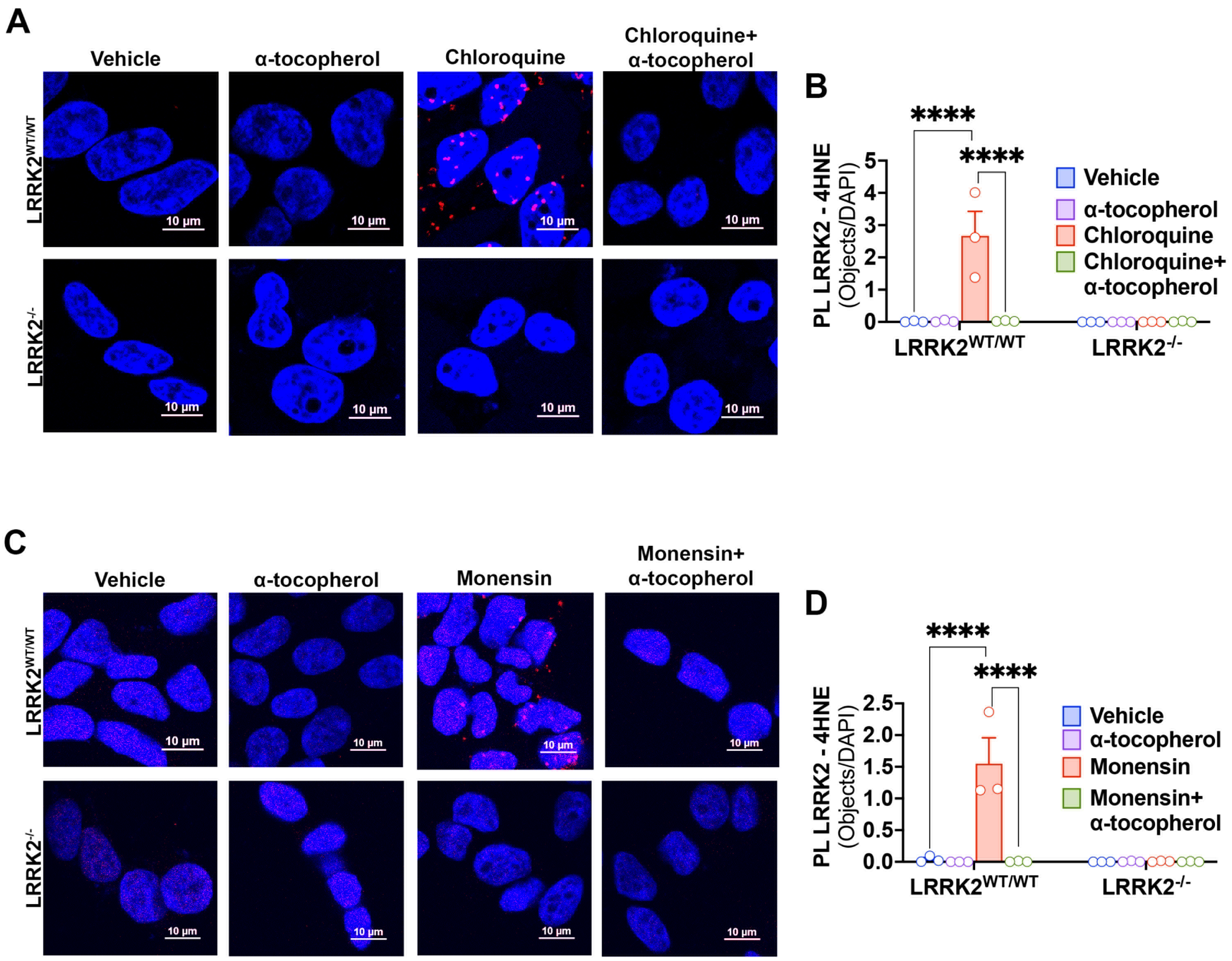


**Fig S10. LRRK2 activating stimuli promote endogenous LRRK2–4-HNE adduct formation**. (**A**) PL between LRRK2 and 4-HNE (PL LRRK2–4HNE) was used as an index of endogenous LRRK2–4-HNE adduct formation in $LRRK2^{WT/WT}$ and $LRRK2^{-/-}$ HEK293 cells. $LRRK2^{WT/WT}$ treated with chloroquine had an increase in PL LRRK2-4HNE (red dots), that was prevented by co-treatment with $\alpha$-tocopherol. (**B**) Quantification of PL LRRK2-4HNE Objects/DAPI in HEK293 cells. Each symbol represents the average number of PL LRRK2-4HNE Objects/DAPI obtained from 100-150 cells per treatment group for each independent experiment; N=3 independent experiments. Statistical testing by two-way ANOVA with post-hoc Tukey correction. (**C**) $LRRK2^{WT/WT}$ HEK293 cells exposed to monensin displayed an increase in PL LRRK2-4HNE signal compared to vehicle. This was prevented by co-treatment with $\alpha$-tocopherol. (**D**) Quantification of PL LRRK2-4HNE Objects/DAPI in HEK293 cells. Each symbol represents the average number of PL LRRK2-4HNE Objects/DAPI obtained from 100-150 cells per treatment group for each of three independent experiments. Statistical testing by two-way ANOVA with post-hoc Tukey correction. **** denotes p<0.0001.

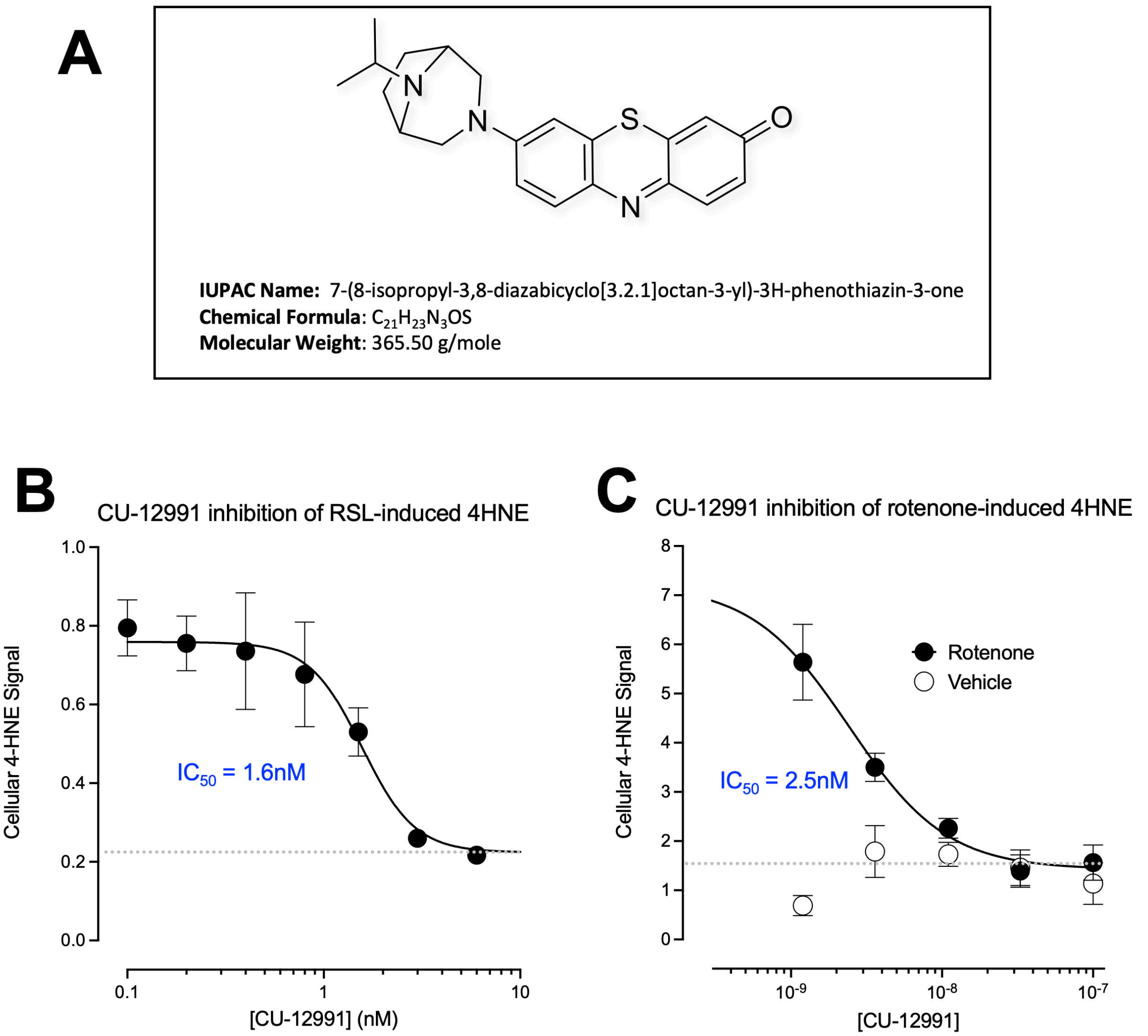


**Fig S11. Structure and potency of CU-12991.** (**A**) Structure of the specific 15-LO inhibitor, CU-12991. (**B**) Dose response curve of cellular 4-HNE signal vs. CU-12991 concentration in $LRRK2^{G2019S}$ fibroblasts treated with the glutathione peroxidase 4 inactivator and ferroptosis inducer, RSL3. Dashed line represents the baseline 4-HNE signal. (**C**) Dose response curve of cellular 4-HNE signal vs. CU-12991 concentration in $LRRK2^{WT/WT}$ RAW264.7 macrophages exposed to 50nM rotenone. Dashed line represents the baseline 4-HNE signal.

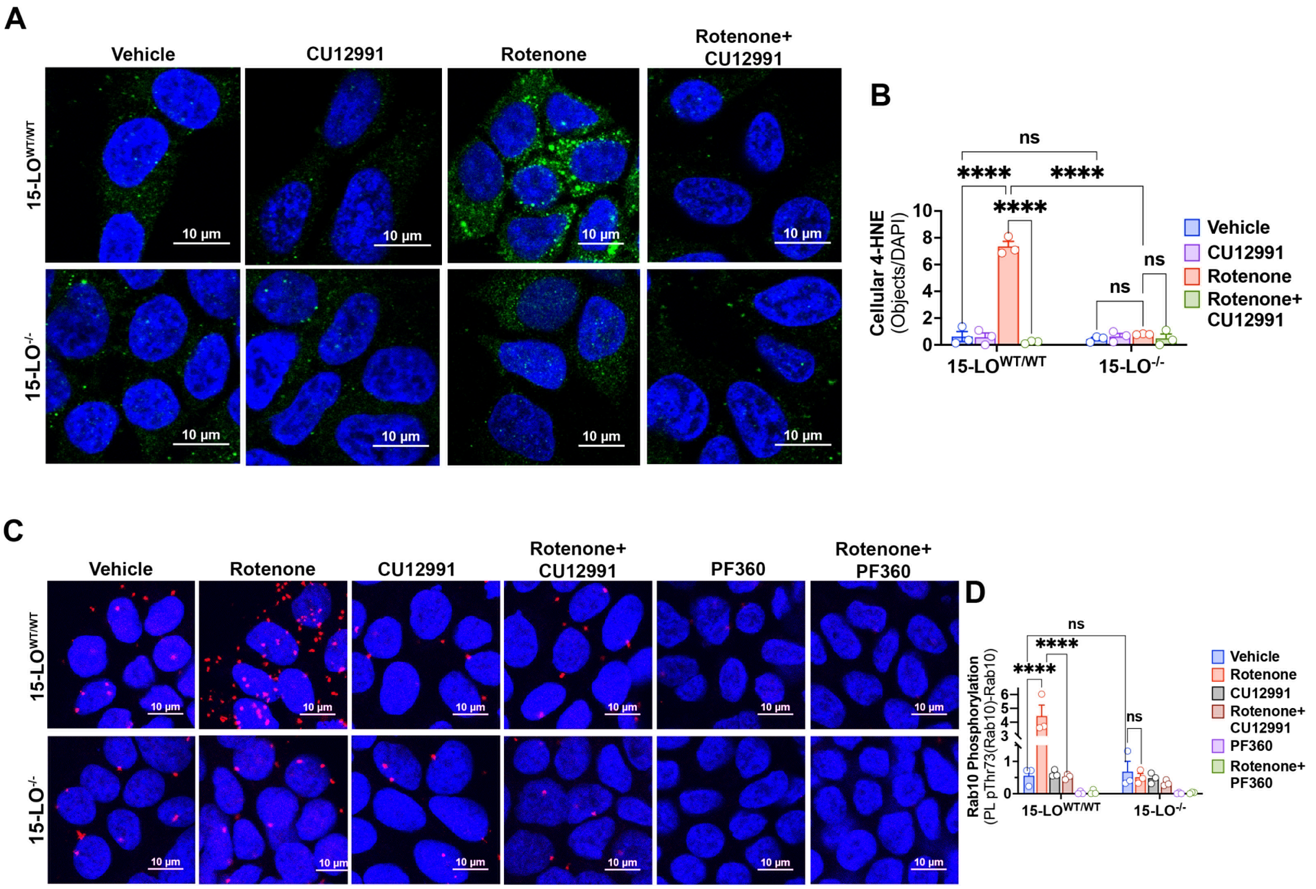


**Fig S12. 15-LO$^{-/-}$ prevents rotenone induced 4-HNE accumulation and Rab10 phosphorylation.** (**A**) Immunofluorescence staining for 4-HNE (green) in 15-LO$^{WT/WT}$ and 15-LO$^{-/-}$ HAP1 cells. Rotenone treatment elicited an increase in 4-HNE signal in 15-LO$^{WT/WT}$, which was prevented by the 15-LO inhibitor, CU-12991. In 15-LO$^{-/-}$ HAP1 cells, rotenone induced 4-HNE accumulation was prevented. (**B**) Quantification of cellular 4-HNE signal. Each symbol represents the average number of 4-HNE objects/DAPI values obtained from 100-150 cells per treatment group per experiment; N=3 independent experiments. Statistical testing by two-way ANOVA with post-hoc Tukey correction. (**C**) PL pT73(Rab10)-Rab10 (red dots) was used to assess rotenone induced Rab10 phosphorylation as a surrogate marker for LRRK2 kinase activity. 15-LO$^{WT/WT}$ cells treated with rotenone had an increase in PL pT73(Rab10)–Rab10 signal (red dots) relative to vehicle, which was prevented by cotreatment with CU-12991 or completely abolished by PF360. Baseline pT73(Rab10)-Rab10 signal was similar between 15-LO$^{-/-}$ and 15-LO$^{WT/WT}$, however rotenone did not result in an increase in PL pT73(Rab10)-Rab10 signal in 15-LO$^{-/-}$. (**D**) Quantification of Rab10 phosphorylation by PL pT73(Rab10)-Rab10 Objects/DAPI. Each symbol represents the average number PL pT73(Rab10)-Rab10 Objects/DAPI obtained from 100-150 cells per treatment group for each of three independent experiments. Statistical testing by two-way ANOVA with post-hoc Tukey correction. **** denotes p<0.001.

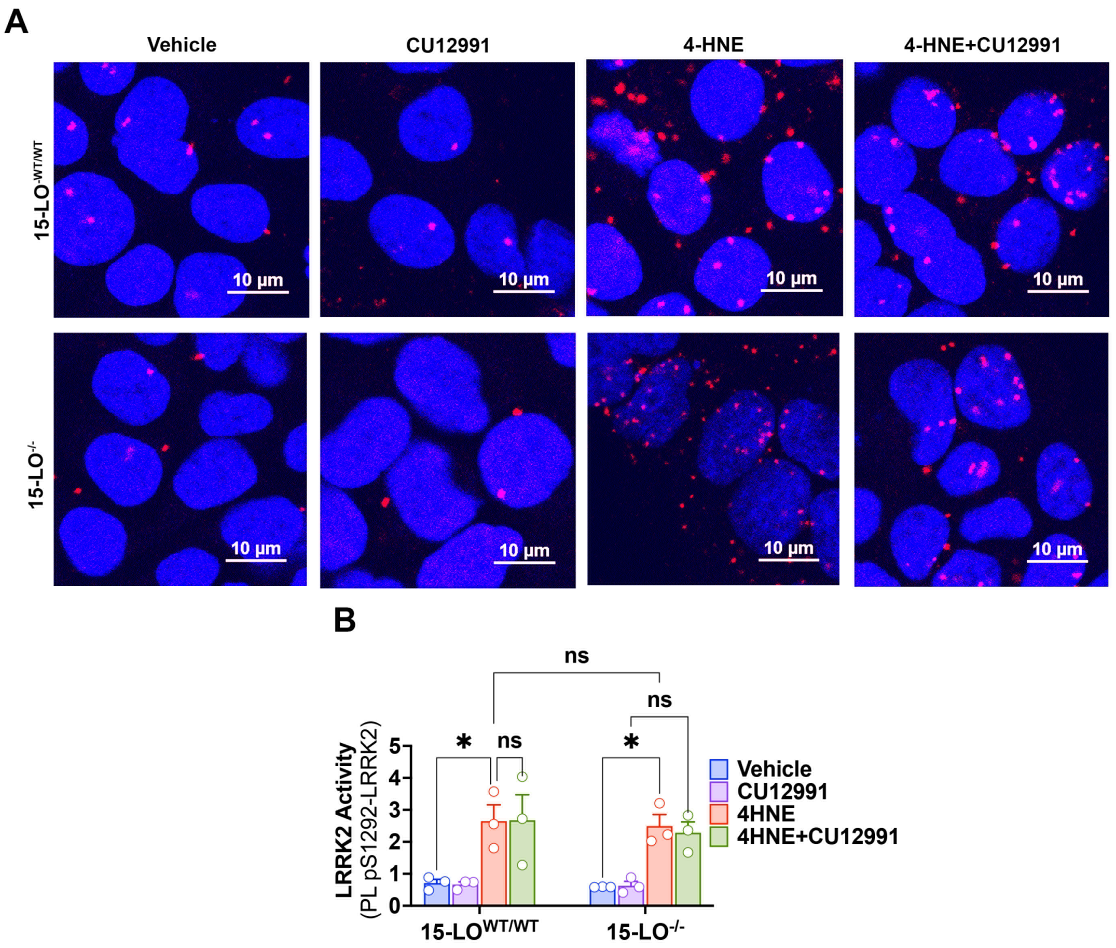


**Fig S13. CU-12991 does not interfere with 4-HNE.** (**A**) 4-HNE induced LRRK2 kinase activity measured by PL pS1292-LRRK2 was assessed in 15-LO$^{WT/WT}$ and 15-LO$^{-/-}$ HAP1 cells in presence or absence of CU-12991. 4-HNE treatment elicited an increase in PL pS1292-LRRK2 in both in 15-LO$^{WT/WT}$ and 15-LO$^{-/-}$ cells. CU-12991 co-treatment had no effect on 4-HNE induced PL pS1292-LRRK2 in 15-LO$^{WT/WT}$ and 15-LO$^{-/-}$ cells. (**B**) Quantification of LRRK2 activity by PL pS1292-LRRK2 Objects/DAPI. Each symbol represents the average number of PL pS1292-LRRK2 objects/DAPI obtained from 100-150 cells per treatment group for each of three independent experiments. Statistical testing by two-way ANOVA with post-hoc Tukey correction. * denotes $p<0.05$.

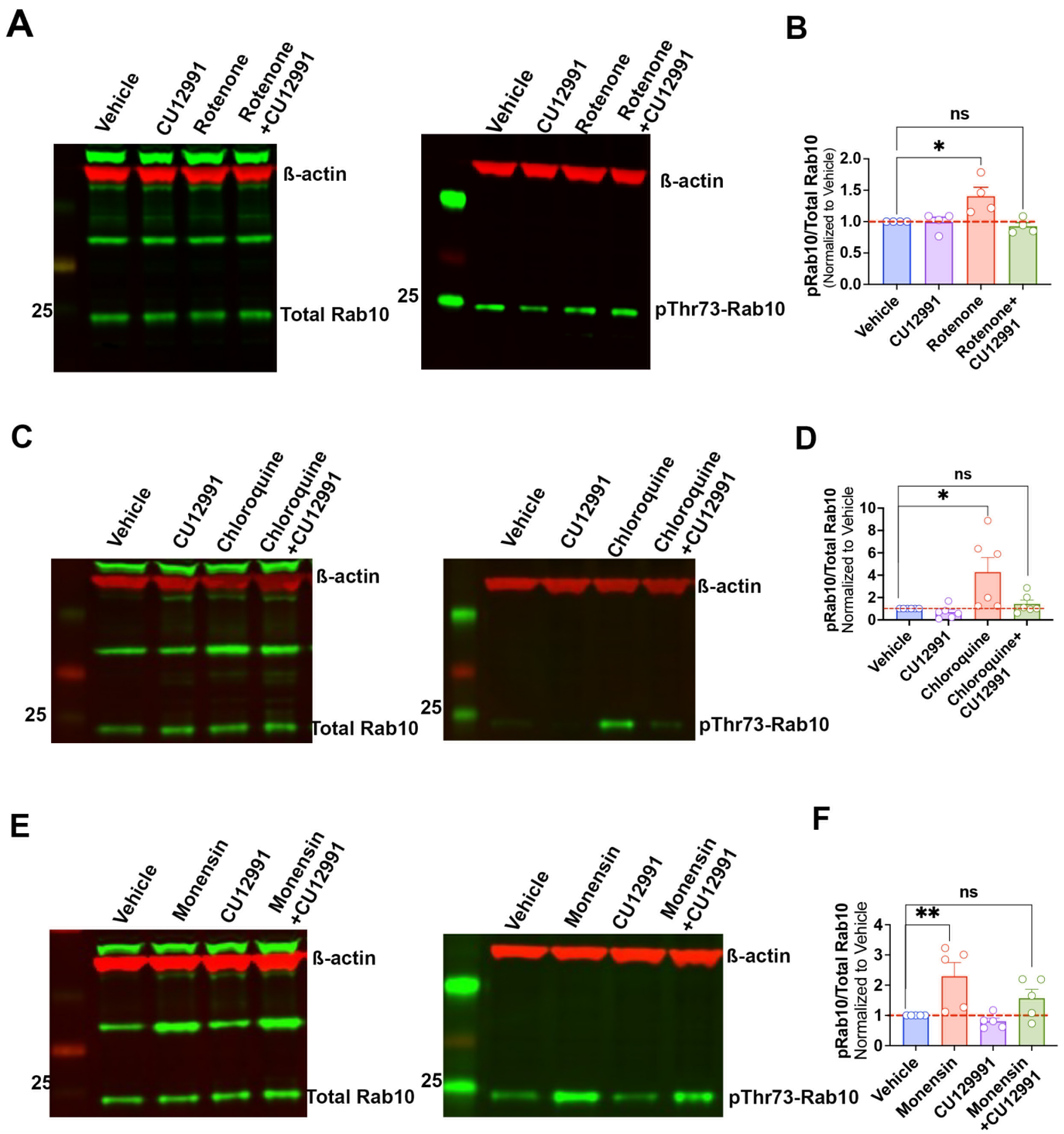


**Fig S14. CU-12991 prevents stimulated Rab10 phosphorylation**. Representative western blots for total Rab10 and pThr73-Rab10 in LRRK2$^{WT/WT}$ RAW264.7 macrophages treated with (**A**) rotenone, (**B**) chloroquine, or (**C**) monensin. (**A**) Rotenone, (**B**) chloroquine, and (**C**) monensin elicited an increase in pT73-Rab10/Total Rab10 compared to vehicle, which was prevented by co-treatment with CU-12991. (**B**) Quantification of rotenone induced pRab10/Total Rab10 normalized to vehicle. Each symbol represents the average of pT73-Rab10/Total Rab10 obtained from two technical replicates per treatment group for each experiment; N=4 independent experiments. Statistical testing by two-way ANOVA with post-hoc Tukey correction. (**D**) Quantification of chloroquine induced pRab10/Total Rab10 normalized to vehicle. Each symbol represents the average of pT73-Rab10/Total Rab10 obtained from two technical replicates per treatment group for each experiment; N=6 independent experiments. Statistical testing by two-way ANOVA with post-hoc Tukey correction. (**F**) Quantification of monensin induced pRab10/Total Rab10 normalized to vehicle. Each symbol represents the average of pT73-Rab10/Total Rab10 obtained from two technical replicates per treatment group for each of five experiments. Statistical testing by two-way ANOVA with Tukey correction. * denotes $p<0.05$; ** denotes $p<0.005$; ns denotes not significant.

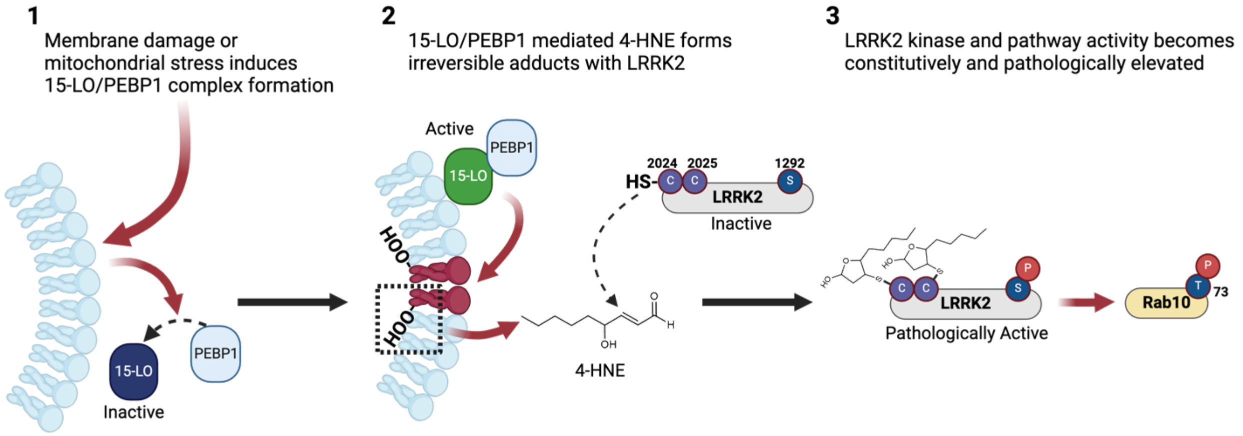


**Fig S15. 4-HNE mediated LRRK2 activation.** (**1**) Under physiological conditions, 15-LO prefers free fatty acids as substrates, not those incorporated into membranes. Under PD-related pathogenic conditions (i.e., mitochondrial stress, lysosomal dyshomeostasis, or vesicular trafficking impairment), the scaffolding protein PEBP1 associates with 15-LO. (**2**) The newly formed PEBP1–15-LO complex allows 15-LO to preferentially hydroperoxidate membrane phospholipids, the first key step in the enzymatic generation of 4-HNE. Sustained 15-LO activation, resulting from these cellular perturbations, leads to the excessive generation and accumulation of 4-HNE at membranes and in the cytoplasm. 4-HNE is a reactive aldehyde that forms adducts with proteins such as LRRK2 and influences their function. (**3**) LRRK2–4-HNE adduct formation results in the hyperactivation of LRRK2 kinase and the LRRK2 pathway, leading to elevated levels of pT73 Rab10. Pharmacological inhibition of 15-LO, 15-LO genetic knock-out, or scavenging the endogenous generation of 4-HNE mitigates pathological LRRK2 kinase and pathway activation. Thus, 15-LO regulates pathological LRRK2 kinase activity through the 15-LO-mediated generation of the lipid hydroperoxidation end-product 4-HNE.

**Supplemental Table 1:** Reagents used for cell culture treatments

| Reagent | Catalog | Vendor | Concentration | Time |
|---|---|---|---|---|
| Rotenone | R8875-1G | Sigma Aldrich | 50nM | 24-hours |
| $H_2O_2$ | 216763 | Sigma Aldrich | 5µM | See time-course Fig. 1 |
| PF360 | | Gift from Pfizer | 1µM | 24-hours |
| **4-HNE** | 31200 | Caymen Chemical | 10µM, 30µM, 100µM | 1 hour |
| 15(S)-HpETE | 44720 | Caymen Chemical | 0.25µM, 2.5µM, 25µM | 1 hour |
| 12(S)-HpETE | 44570 | Caymen Chemical | 0.25µM, 2.5µM, 25µM | 1 hour |
| Chloroquine | C6628 | Sigma Aldrich | 100µM | 3-hours or 1 hour (4-HNE assay) |
| Monensin | M5273 | Sigma Aldrich | 10µM | 4 hours or 1 hour (4-HNE assay) |
| N-acetyl-L-cysteine (NAC) | A7250 | Sigma Aldrich | 250µM | 24-hours; 2-hour pretreatment for lipid experiments |
| α-tocopherol | T3251 | Sigma Aldrich | 100µM | 24-hours |
| CU-12991 | | Acurex Biosciences | 20nM | 24-hours |

**Supplemental Table 2**: Lymphoblastoid cell line information

| Cell line ID | LRRK2 Mutation | Clinical Status | Sex | Age at sampling |
|---|---|---|---|---|
| ND00312 | None detected | Unaffected | M | 72 |
| ND01277 | None detected | Unaffected | F | 68 |
| ND02379 | None detected | Unaffected | M | 51 |
| ND02599 | None detected | Unaffected | M | 68 |
| ND00075 | G2019S | Parkinson's disease (PD) | M | 50 |
| ND14317 | G2019S | PD | M | 53 |
| ND03000 | G2019S | PD | F | 68 |
| ND00045 | G2019S | PD | F | 60 |
| ND03899 | None detected | idiopathic PD (iPD) | M | 53 |
| ND01180 | None detected | iPD | M | 50 |
| ND01979 | None detected | iPD | F | 75 |
| ND01610 | None detected | iPD | F | 68 |

**Supplemental Table 3**: CRISPR/Cas9 reagents

| | |
|---|---|
| **Exon 41 Forward** | **5'-TCCAAAAATTGGGTCTTTGC-3'** |
| **Exon 41 Reverse** | **5'-CACAATGTGATGCTTGCATTT-3'** |
| **LRRK2 Guide RNA** | **5'-CTCAGTACTGCTGTAGAATG-3'** |
| **C2024A repair template** | **5'-ACACTG TATCCCAATGCTGCCATCATTGCAAAGATTGCTGACTACGGCATTGCTCA GTACGCCTGTAGAATGGGGATAAAAACATCAGAGGGCACACCAGgtaggtgatcaggtctgtct-3'** |
| **C2025A repair template** | **5'-ACACTG TATCCCAATGCTGCCATCATTGCAAAGATTGCTGACTACGGCATTGCTCA GTACTGCGCTAGAATGGGGATAAAAACATCAGAGGGCACACCAGgtaggtgatcaggtctgtct-3'** |
| **C2024A/ C2025A repair template** | **5'-ACACTG TATCCCAATGCTGCCATCATTGCAAAGATTGCTGACTACGGCATTGCTCA GTACGCCGCTAGAATGGGGATAAAAACATCAGAGGGCACACCAGgtaggtgatcaggtetgtct-3’** |

**Supplemental Table 4**: List of antibodies using for immunofluorescence, proximity ligation assay and western blot

| Antibody | Catalog | Vendor | Dilution |
|---|---|---|---|
| Rabbit pS1292-LRRK2 | ab203181 | Abcam | 1:1,000 (PL pS1292-LRRK2) |
| Mouse LRRK2 (N241A) | 75-253 | Antibodies Inc. | 1:1,000 (PL pS1292-LRRK2 and western blot)<br>1:500 (PL LRRK2-4HNE) |
| Rabbit pT73-Rab10 | ab230261 | Abcam | 1:1,000 (western blot) |
| Rabbit pT73-Rab10 | ab241060 | | 1:1,000 (PL pT73(Rab10) – Rab10) |
| Rabbit Total Rab10 | ab237703 | Abcam | 1:2,000 (western blot) |
| Mouse Total Rab10 | ab104859 | Abcam | 1:1,000 (PL pRab10 – Rab10) |
| Rabbit PEBP/RKIP | 13006S | Cell Signaling | 1:500 (IF and PL PEBP1 – 15-LO cells )<br>1:250 (PL PEBP1 – 15-LO tissue) |
| Mouse 15-Lipoxygenase | sc-133085 | Santa Cruz | 1:1,000 (IF and PL PEBP1 – 15-LO cells)<br>1:250 (PL PEBP1 – 15-LO tissue) |
| Mouse β-actin | NBP1-47423 | Novus Biologicals | 1:5,000 (western blot) |
| Rabbit 4-HNE | ab46545 | Abcam | 1:1,000 (IF)<br>1:500 (PL LRRK2-4HNE, cells and tissue) |
| Sheep tyrosine hydroxylase | AB1542 | Millipore Sigma | 1:2,000 (IHC) |
| IRDye 800CW Donkey anti-Mouse IgG Secondary Antibody | 926-32212 | LiCor | 1:5,000 |
| IRDye 680RD Donkey anti-Mouse IgG Secondary Antibody | 926-68072 | LiCor | 1:5,000 |

**Supplemental Table 4**: List of antibodies using for immunofluorescence, proximity ligation assay and western blot

| Antibody | Catalog | Vendor | Dilution |
|---|---|---|---|
| Rabbit pS1292-LRRK2 | ab203181 | Abcam | 1:1,000 (PL pS1292-LRRK2) |
| Mouse LRRK2 (N241A) | 75-253 | Antibodies Inc. | 1:1,000 (PL pS1292-LRRK2 and western blot)<br>1:500 (PL LRRK2-4HNE) |
| Rabbit pT73-Rab10 | ab230261 | Abcam | 1:1,000 (western blot) |
| Rabbit pT73-Rab10 | ab241060 | | 1:1,000 (PL pT73(Rab10) – Rab10) |
| Rabbit Total Rab10 | ab237703 | Abcam | 1:2,000 (western blot) |
| Mouse Total Rab10 | ab104859 | Abcam | 1:1,000 (PL pRab10 – Rab10) |
| Rabbit PEBP1/RKIP | 13006S | Cell Signaling | 1:500 (IF and PL PEBP1 – 15-LO cells )<br>1:250 (PL PEBP1 – 15-LO tissue) |
| Mouse 15-Lipoxygenase | sc-133085 | Santa Cruz | 1:1,000 (IF and PL PEBP1 – 15-LO cells)<br>1:250 (PL PEBP1 – 15-LO tissue) |
| Mouse β-actin | NBP1-47423 | Novus Biologicals | 1:5,000 (western blot) |
| Rabbit 4-HNE | ab46545 | Abcam | 1:1,000 (IF)<br>1:500 (PL LRRK2-4HNE, cells and tissue) |
| Sheep tyrosine hydroxylase | AB1542 | Millipore Sigma | 1:2,000 (IHC) |
| IRDye 800CW Donkey anti-Mouse IgG Secondary Antibody | 926-32212 | LiCor | 1:5,000 |
| IRDye 680RD Donkey anti-Mouse IgG Secondary Antibody | 926-68072 | LiCor | 1:5,000 |
